# Dynamic regulation of inter-organelle communication by ubiquitylation controls skeletal muscle development and disease onset

**DOI:** 10.1101/2022.07.21.501000

**Authors:** Arian Mansur, Remi Joseph, Pierre Michael Jean-Beltran, Namrata D. Udeshi, Cadence Pearce, Hanjie Jiang, Reina Iwase, Elyshia McNamara, Jeff Widrick, Claudio Perez, Gianina Ravenscroft, Philip A Cole, Steven A Carr, Vandana A Gupta

## Abstract

Ubiquitin-proteasome system (UPS) dysfunction is associated with the pathology of a wide range of human diseases including myopathies and muscular atrophy. However, the mechanistic understanding of specific components on the regulation of protein turnover during development and disease progression in skeletal muscle is unclear. Mutations in *KLHL40*, an E3 ubiquitin ligase cullin3 (CUL3) substrate-specific adapter protein result in a severe form of congenital nemaline myopathy, but the events that initiate the pathology and the mechanism through which it becomes pervasive, remains poorly understood. To characterize the KLHL40-regulated ubiquitin modified proteome during skeletal muscle development and disease onset, we used global, quantitative mass spectrometry-based ubiquitylome and global proteome analyses of *klhl40* mutant zebrafish during disease progression. Global proteomics during skeletal muscle development revealed extensive remodeling of functional modules linked with sarcomere formation, energy and biosynthetic metabolic processes and vesicle trafficking. Combined analysis of *klh40* mutant muscle proteome and ubiquitylome identified thin filament proteins, metabolic enzymes and ER-Golgi vesicle trafficking pathway proteins regulated by ubiquitylation during muscle development. Our studies identified a role for KLHL40 as a negative regulator of ER-Golgi anterograde trafficking through ubiquitin-mediated protein degradation of secretion associated Ras related GTPase1a (Sar1a). In KLHL40 deficient muscle, defects in ER exit site vesicle formation alter Golgi compartment and downstream transport of extracellular cargo proteins, resulting in structural and functional abnormalities. Our work reveals that the muscle proteome is dynamically fine-tuned by ubiquitylation to regulate skeletal muscle development and uncovers new disease mechanisms for therapeutic development in patients.

**Graphical Abstract:** 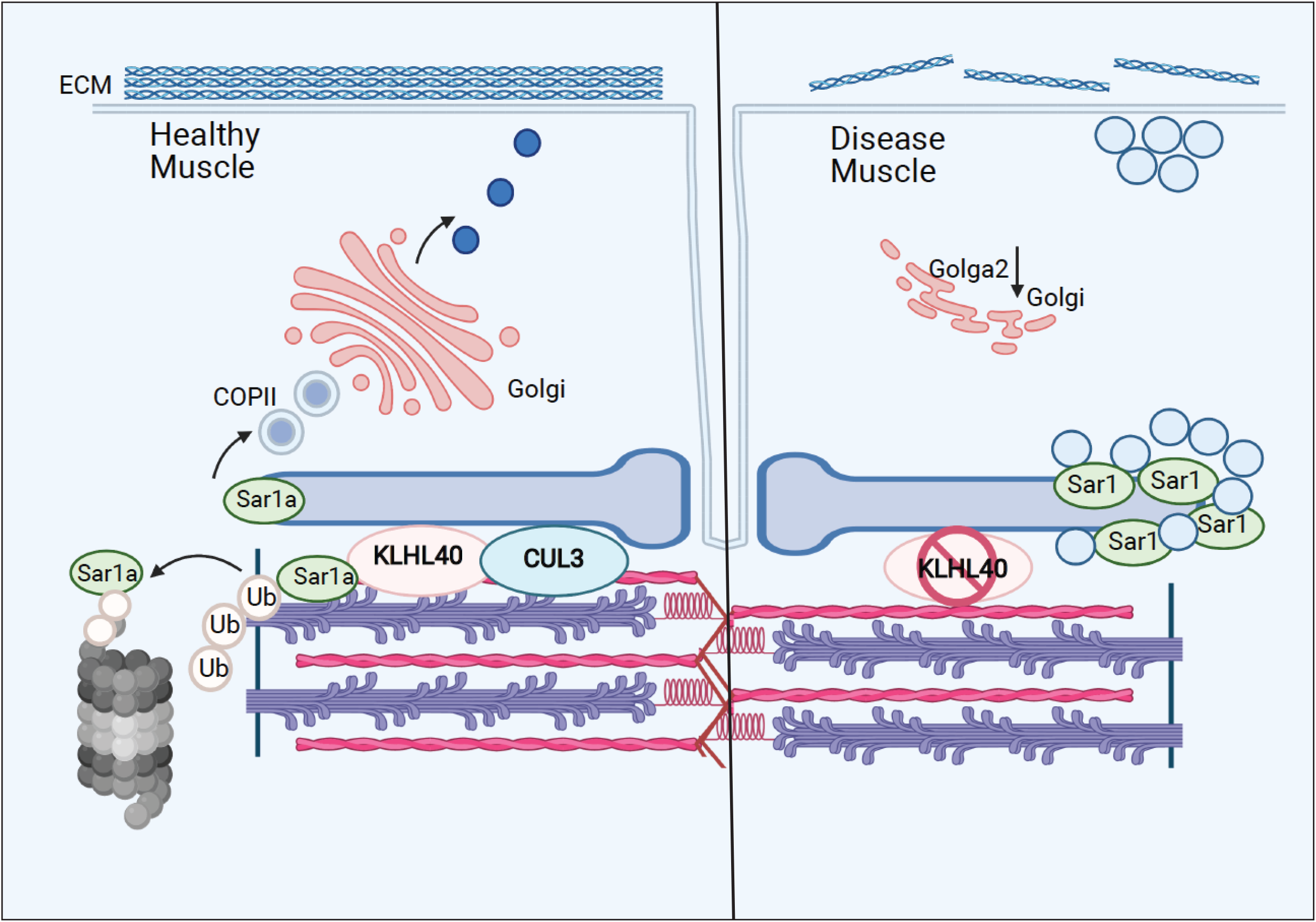

## Introduction

Fetal akinesia, arthrogryposis and severe congenital myopathies are heterogeneous conditions of reduced fetal movement usually presenting at birth (1, 2). More than 50% of all causes of fetal akinesia are of neuromuscular origin, involving all points along the neuromuscular axis (motor neurons, peripheral nerves, neuromuscular junction, and the skeletal muscle regulatory and contractile apparatus) (3-7). These diseases exhibit a high clinical heterogeneity with a severe congenital onset with fetal akinesia to milder forms often with a late childhood or an adult-onset. At least 30 causative genes have been identified in these conditions (8, 9). However, the origin and temporal ordering of molecular events that drive the disease pathology remains poorly understood.

Skeletal muscle is made up of myofibers extremely specialized for contraction. To achieve this function, each myofiber contains myofibrils, which consist of a repetition of sarcomeres. Subsequent to myoblast fusion, sarcomeres are assembled through the interaction of protein complexes that form complex supramolecular structures to form functional myofibers. This requires precisely controlled dynamic turn-over of proteins without perturbing the structure of assembling sarcomeres. The ubiquitin proteasome system regulates the relative abundance and functional modifications of proteins during multiple stages of myogenesis (10-12). The importance of this process in skeletal muscle development is identified by human diseases where mutations in genes regulating ubiquitination and protein turnover processes result in sarcomeric disarray and functional deficits (3, 13-15).

Sarcomeres are present in close proximity to the triad system that is formed of T-tubules and sarcoplasmic reticulum, a modified endoplasmic compartment. In addition, different mitochondrial populations are also present in close juxtaposition to the sarcomere (16). Mutations or deletions of sarcomeric genes affect the structure and function of surrounding organelles and similarly, defects in other organelles in myofibers also affect sarcomere structure and function (17-20). This suggests that the sarcomere and surrounding organelles act as interconnected hubs that engage in extensive communication during skeletal muscle development and maintenance. Despite this, mechanistic insight into inter-organelle communication and regulation of this process in skeletal muscle development remains largely unknown. Finally, how this communication is perturbed in disease states and contributes to disease pathology is not clear.

Here, we have identified that inter-organelle communication is critical for vesicle trafficking and skeletal muscle development by ubiquitination signaling in the sarcomere. We performed global proteomic and ubiquitylome profiling of skeletal muscle during development and disease progression in a Klhl40 deficient zebrafish model of congenital nemaline myopathy. We identified that sarcomeric KLHL40 acts as a negative regulator of membrane vesicle trafficking through ubiquitylation and subsequent protein degradation of Secretion associated Ras-related GTPase1a (SAR1a). In the absence of this negative feedback mechanism in KLHL40 deficiency, SAR1a is abnormally localized to the ER and contributes to membrane tubulation defects and disruption of the trafficking of ECM proteins. Our work demonstrates that inter-organelle communication between sarcomeric and endomembrane compartments, is dynamically regulated by ubiquitylation and is critical for skeletal muscle development and defects in this process underlie pathology in skeletal muscle diseases.

## Results

### KLHL40 is required for skeletal muscle development

KLHL40 deficiency in humans results in a severe form of nemaline myopathy associated with neonatal lethality. Deletion of *Klhl40* in mice also results in extensive structural damage in myofibers and neonatal lethality two-three weeks after birth. As zebrafish grow *ex vivo*, skeletal muscle development and disease progression can be visualized in the context of a living organism. We generated loss-of-function *klhl40* alleles in zebrafish using the CRISPR/Cas9 gene editing tool (Figure 1A-B). The human orthologue of *KLHL40* gene is duplicated in zebrafish as *klhl40a* and *klhl40b* and mapped on chromosome 2 and chromosome 24, respectively. *klhl40a* alleles created include *klhl40a*^bwg200^ with insertion of one base (c.250_251insA; p.Val84Aspfs*36) and *klhl40a*^bwg201^ with a two base pair deletion in exon 1 (c.251_252insTC;p.Val84Asnfs*36). These alleles result in out of frame mutations and are predicted to result in truncating proteins in the N terminal BTB domain of the Klhl40a protein. For *klhl40b*, a CRISPR edited allele (*klhl40b*^bwg202^) had insertion of one base (c.674_675insC; p.Arg225Profs*14) in exon 1 (Figure 1B). *klhl40b*^bwg202^ allele is predicted to result in an out of frame mutation and truncation of the Klhl40b protein in the BACK domain (Figure 1A). As *klhl40a*^bwg200^ and *klhl40b*^bwg202^ alleles were predicted to produce the smallest truncated Klhl40 proteins. The rest of the analyses presented in this work were performed on these fish lines obtained after F3 generation and referred as *klhl40a* and *klhl40b* in rest of this work. The effect of different mutations on *klhl40* mRNA levels was evaluated by RT-PCR (Figure S1A and B). Both *klhl40a* and *klhl40b* alleles exhibited similar *klhl40a* and *klhl40b* mRNA levels, respectively in comparison to +/+ siblings. To evaluate the effect of these mutations on the klhl40 protein, Western blot was performed on the skeletal muscle extracts obtained from *klhl40a* and *klhl40b* mutant fish. Western blot analysis with a KLHL40 antibody that recognizes both Klhl40a and Klhl40b proteins revealed about 50% decrease in Klhl40 protein levels in both alleles in comparison to control and complete absence of Klhl40 protein in *klhl40a*/*klhl40b*, double knockout fish (Figure S1C and D). This suggests that *klhl40a* and *klhl40b* alleles result in loss of Klhl40 protein in the mutant zebrafish.

**Figure 1.**
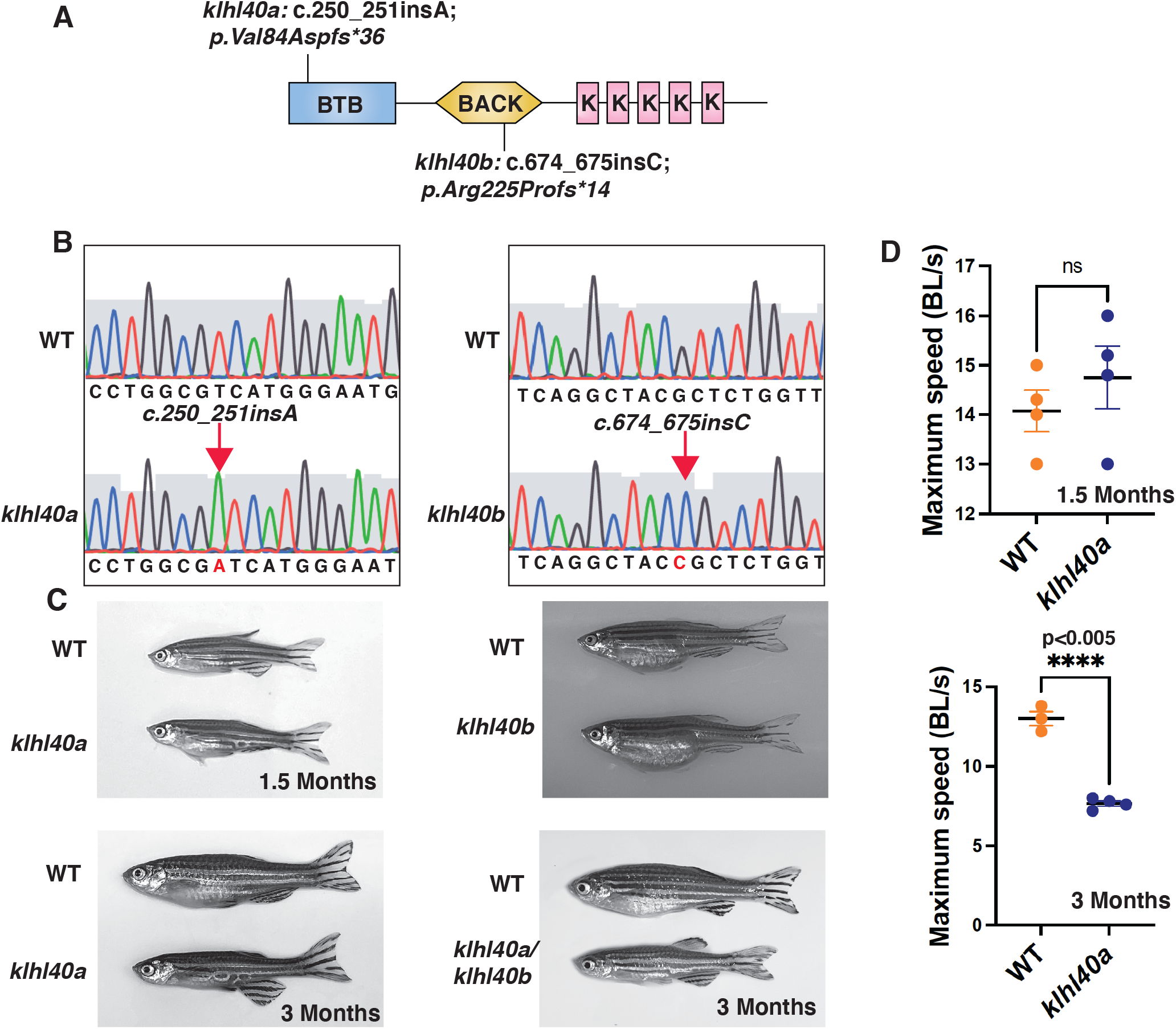
KLHL40 is essential for vertebrate skeletal muscle development. (A) Schematic diagram depicting the position of CRISPR-mediated mutant alleles and truncated proteins on Kelch protein domain in *klhl40a*^*bwg200*^ and *klhl40b*^*bwg202*^ knockout zebrafish. CRISPR-induced mutations in *klhl40a*^*bwg200*^ and *klhl40b*^*bwg202*^ knockout zebrafish result in premature termination codons in the BTB domain and BACK domain coding exons, respectively. (B) Sanger sequencing pherograms for control and *klhl40a* ^*bwg200*^ and *klhl41b* ^*bwg202*^ mutant zebrafish with an insertion of A in *klhl40a*^*bwg200*^ and an insertion of C in *klhl40b*^*bwg202*^ coding regions. (C) Lateral view of the juvenile and adult zebrafish. *klhl41a* mutant zebrafish develop myopathy from transition to juvenile (1.5 months old) to the onset of adult stage (3 months old) and exhibit reduced body length and body diameter. No obvious skeletal muscle phenotype is observed in *klhl41a* allele in comparison to control (+/+) siblings. *klhl40a/ klhl40b* double mutant fish exhibit similar skeletal phenotype as observed in *klhl40a* allele. (D) Endurance swimming behavior of *klhl40a* allele at juvenile state (1.5 months) and adult stage (3 months) (n=7-8). Data are mean±S.E.M (unpaired t-test, parametric) for each experiment.

*klhl40a* and *klhl40b* mutant embryonic (2dpf) and larval fish (5dpf) did not exhibit any gross morphological defects in any of the mutants examined. Previous studies have shown that knockdown of *klhl40* by morpholino results in myopathy in zebrafish embryos (3). The discrepancy between the morphants and mutants could be due to genetic compensatory mechanisms by other Kelch protein coding genes or other modifier genes as described by several studies (21, 22). During the developmental transition from juvenile (1.5 months) to adult stage (3.0 months), the *klhl40a* mutants develop a myopathic phenotype with reduced body size whereas *klhl40b* were phenotypically indistinguishable from +/+ control siblings at this age (Figure 1C). *klhl40a*/*klhl40b* double mutants appeared to be phenotypically similar to *klhl40a* fish suggesting that duplicated *klhl40b* is redundant for normal muscle growth. To identify any defects in skeletal muscle function, the swimming performance of *klhl40a* mutants and +/+ siblings were analyzed by the flume tunnel assay to obtain maximum swimming speed (*U*_*max*_) (Figure 1D) (23). No differences in the *U*_*max*_ values were observed between control and *klhl40a* mutants at juvenile stage (1.5 months). The *U*_*max*_ values showed a significant decrease in *U*_*max*_ in *klhl40a* mutants compared to +/+ control siblings at the adult stage (3 months), indicating reduced endurance capacity of the klhl40a deficient fish. As *klhl40a* mutant fish at 1.5 months are phenotypically and functionally similar to control fish, this age group was termed as “pre-symptomatic stage” whereas *klhl40a* mutant fish at 3.0 months was termed as “symptomatic stage” due to myopathic features. Control fish survived to 24 months of age whereas most of the *klhl40a* mutant fish died between 9-12 months. These data show that loss of *klhl40a* leads to a myopathic phenotype as observed in patients with *KLHL40* variants.

### KLHL40 plays pleiotropic roles in regulating skeletal muscle structure

KLHL40 deficient muscle in patients exhibit extensive myofiber damage and extensive sarcomeric disarray in many myofibers. As patient muscle biopsies are typically collected after the disease diagnosis, in most cases disease processes are already established. This eludes the understanding of series of pathological changes that resulted in the extensive muscle damage. Therefore, to understand how KLHL40 deficiency affects skeletal muscle structure during disease onset and progression, ultra-structure was evaluated by transmission electron microscopy (TEM) in both juvenile (pre-symptomatic) and adult stages (symptomatic) in control and *klhl40a* mutant fish. (Figure 2). While no significant ultrastructural changes were observed during the juvenile stage, both sarcomere width (*w*) and height (*h*) were significantly reduced in the *klhl40a* in comparison to +/+ controls at the adult stage (Figure 2A-B). Mutant muscle also displayed accumulation of vesicles (10-100 nm) in the close proximity to intermyofibrillar SR-ER region (Figure 2C-D). Mitochondria in the skeletal muscle of the *klhl40a* mutant muscle displayed an electron dense-matrix in comparison to the controls suggesting KLHL40 deficiency may also trigger mitochondrial damage (Figure 2C-D). Mutant muscle also exhibited abnormalities in the extracellular matrix (ECM) structure in the mutant muscle (Figure 2E-F). In comparison to the controls, mutant muscle exhibited large gaps in the adjacent myofibers in the ECM region. Interestingly, the Golgi complex was fragmented in skeletal muscle of *klhl40a* in comparison to the well-organized Golgi complex in +/+ controls (Figure 2G-H). In addition, there is an accumulation of small vesicles (30-50 nm) surrounding the damaged Golgi complex in the mutant muscles that appear to be similar in size and electron density to terminal cisternae of the sarcoplasmic reticulum. In summary, these data indicate that KLHL40 is required to regulate sarcomere size, intracellular membrane homeostasis and ECM stability in skeletal muscle.

**Figure 2.**
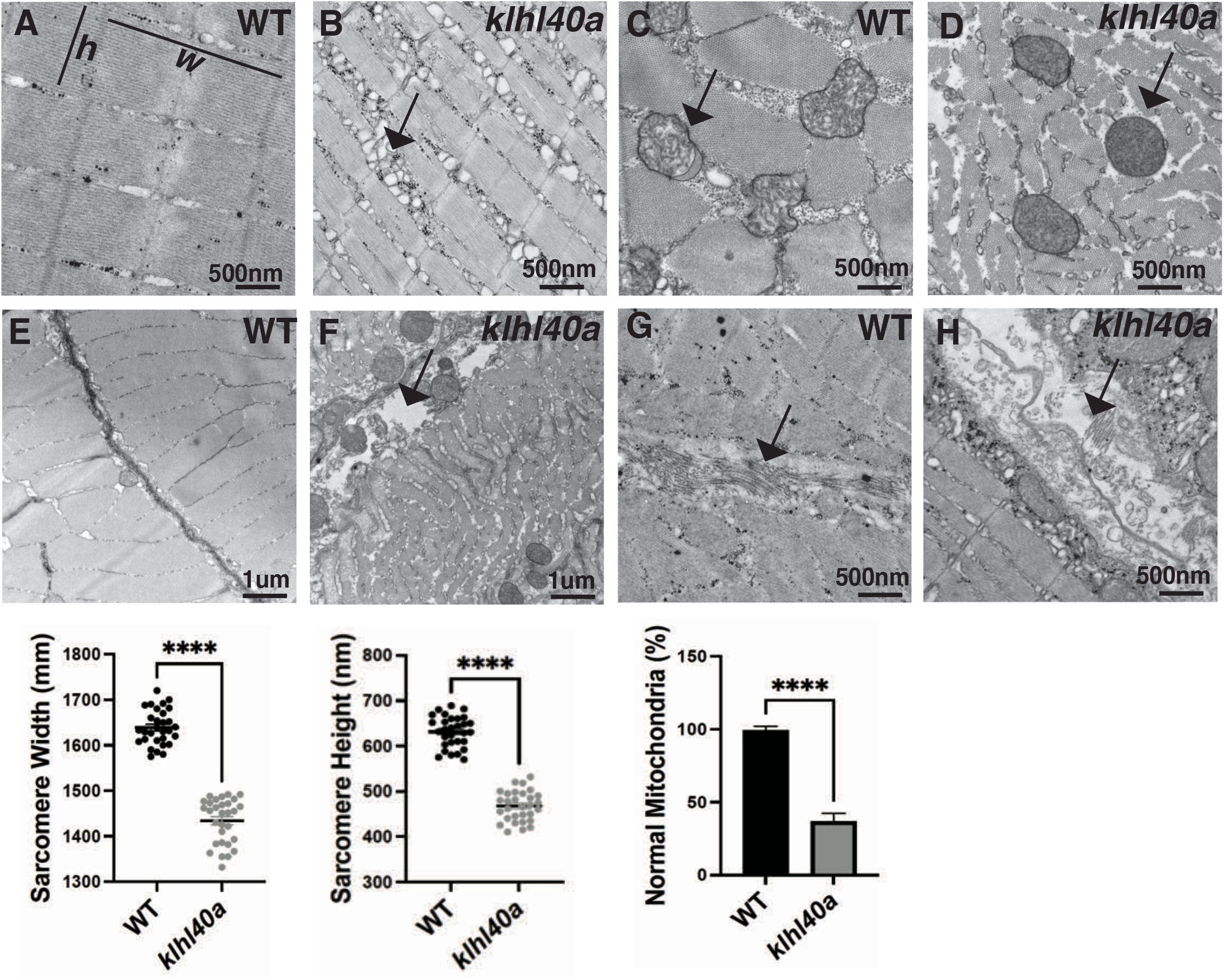
*klhl40a* allele display reduced sarcomere size and abnormal membrane organelles in skeletal muscle. Transmission electron microscopy (TEM) showing ultrastructure of control (+/+) and *klhl40a KO* in 3 months animals. (A-B) Longitudinal muscle section of control and *klhl40a KO* mutant muscle showing accumulation of vesicles in SR-ER region (arrow) and reduced sarcomere width (*w*) and height (*h*). (C-D) Cross-section view showing mitochondrial in *klhl40a KO* mutant muscle contain electron dense matrix (arrow) in comparison to control muscle (normal mitochondria). (E-F) Longitudinal view of skeletal muscle showing structural damage to the extracellular matrix (arrow) in *klhl40a KO* mutant in comparison to the control. (G-H) Longitudinal view of skeletal muscle demonstrating organized Golgi apparatus (arrow) that is fragmented in the *klhl40a KO* mutant muscle. Electron microscopy was performed in three different control and *klhl40a KO* mutant fish. N=150-175 sarcomeres were analyzed in each sample for quantification. N= 75-100 mitochondria were analyzed in each sample for quantification. Data are mean ± s.e.m; with one way analysis of variance (ANOVA) and Tukey’s HSD test (****p<0.001).

### Dynamic remodeling of the proteome during skeletal muscle development and disease progression in nemaline myopathy

KLHL40 is a substrate-specific adaptor of CUL3 E3 ubiquitin ligase and KLHL40-CUL3 ubiquitin ligase complex has previously been shown to stabilize the sarcomeric thin filament proteins such as leimodin3 (LMOD3) and nebulin (NEB) by ubiquitylation through *in vitro* studies (24), however, *in vivo* relevance remains unknown. Protein complexes are changed dynamically during development to meet the constantly changing demands of differentiating cells. Subtle changes in proteins may have significant effect on downstream processes can be hard to identified and require highly quantitative *in vivo* approaches. Identification low abundance proteins with critical roles may pose to be a challenging issue. These issues become particularly significant in human diseases as disease processes are mostly investigated during the pathological states when atrophic processes are prevalent but our understanding can benefit by analyzing disease trajectories from a pre-symptomatic state to clinically symptomatic states.

To comprehensively quantify proteome remodeling during skeletal muscle growth, disease onset and progression, global proteomic changes in skeletal muscle from control (+/+) and *klhl40a* zebrafish at pre-symptomatic (1.5 months) and symptomatic (3 months) stages of disease progression were analyzed. Deep-scale quantitative liquid chromatography tandem mass-spectrometry (LC-MS/MS) based proteomics was performed in skeletal muscle in these fish (Figure 3) (Table S1). Muscle samples from wild-type (WT) and *klhl40a* mutant (KO) were collected and analyzed in quadruplicate using tandem mass tags (TMT) for multiplexing and quantification (Figure 3A) (25). In parallel, we performed deep ubiquitylation profiling via enrichment of the lysine di-glycine remnant (KGG) from ubiquitin trypsinization from the exact same tissues (26). This allows determination of the contribution of ubiquitylation in the remodeling of skeletal muscle proteome during muscle development and disease onset by CUL3 E3 ubiquitin ligase-KLHL40 complex (Table S2) (Figure 3A).

**Figure 3.**
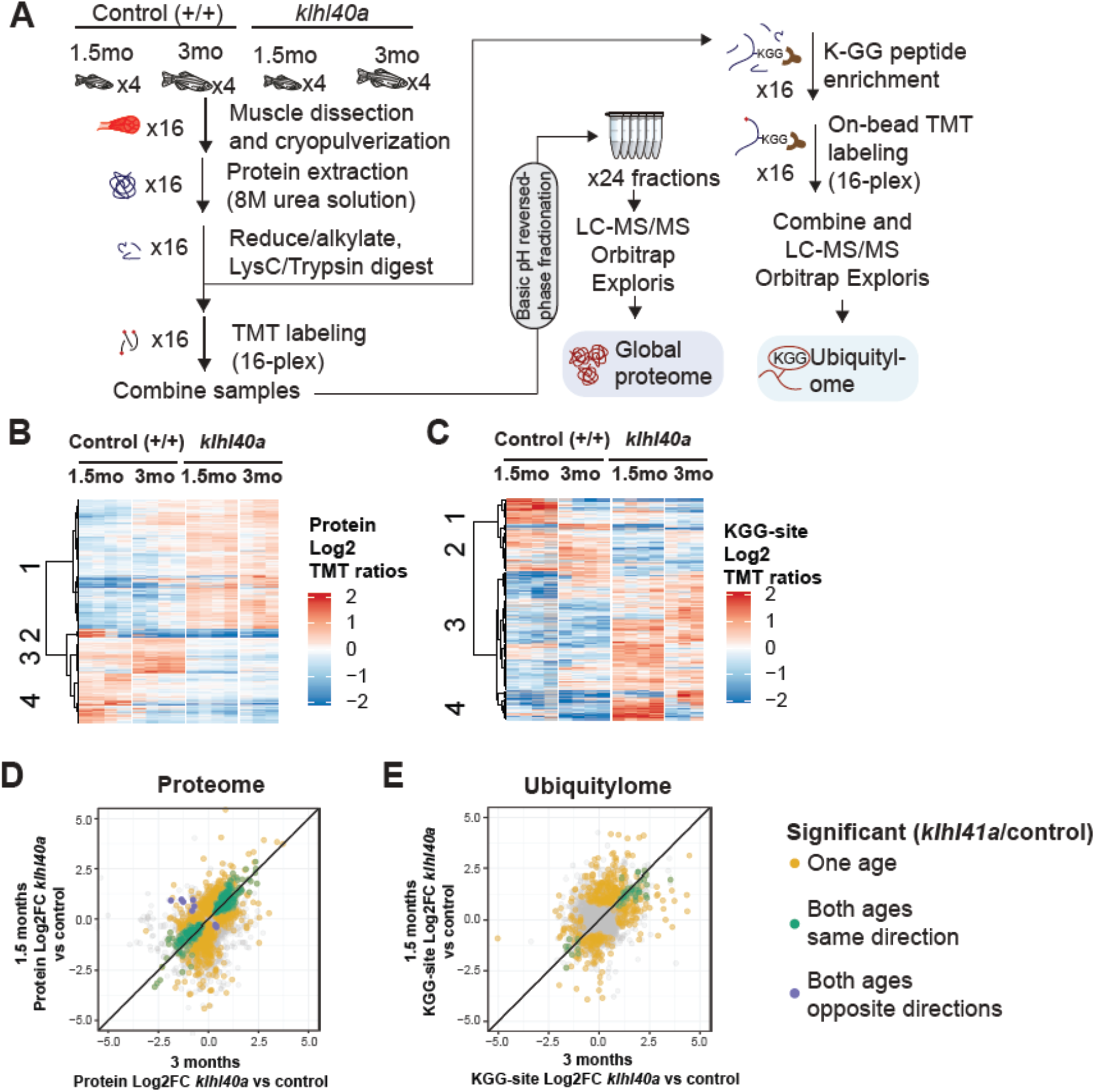
Proteome and ubiquitylome disruption by KLHL40 deficiency. (A) Experimental workflow for proteome and ubiquitylome quantification in *klhl40a* allele. (B) Heatmap showing protein abundances (log2 TMT ratios) across experimental samples. Only proteins with a significant differential response between control and *klhl40a KO* samples are shown (adj. p-val < 0.05). Proteins (rows) were clustered to show abundance patterns across experimental groups. (C) Heatmap showing ubiquitin sites following trypsin digestion (KGG)-site abundances (log2 TMT ratios) across experimental samples. Only proteins with a significant differential response between control and *klhl40a KO* samples are shown (adj. p-val < 0.05). KGG-sites (rows) were clustered to show abundance patterns across experimental groups. (D) Correlation of protein response to *klhl40a KO* across juvenile (1.5 months) and adult animals (3 months). Plots show log2 fold-changes for proteins quantified at both ages. Proteins are colored if they show differential abundance (adj. p-val < 0.05) at one age only (yellow), both ages with same direction (green), and both ages with opposite directions (purple). (E) Correlation of ubiquitylome response to *klhl40a KO* across juvenile (1.5 months) and adult animals (3 months). Plots show log2 fold-changes for KGG-sites quantified at both ages. KGG-sites are colored if they show differential abundance (adj. p-val < 0.05) at one age only (yellow), both ages with same direction (green), and both ages with opposite directions (purple).

A total of 8,268 proteins were quantified across these 16 samples, and PCA showed grouping of the different replicates from each age and genotype experimental group (Figure S2). To investigate changes in the proteome that are primarily regulated through the transcriptome during the disease state, we integrated our proteomic data with RNA sequencing (RNA-seq) results obtained on the same tissue samples (3 months) (Table S3). The proteome-transcriptome correlation analyses revealed a high degree of discordance (78%) between transcript-protein pairs (Figure S3). This reflects extensive post-translational regulation of skeletal muscle development *in vivo* and therefore, we focused on the proteome dataset. Proteins with a significant differential response between control and mutant samples at either stage (moderated t-test, adjusted *p*-value <0.05) were clustered using hierarchical clustering to reveal proteome abundance patterns across experimental groups (Figure 3B). This revealed 4 distinct clusters defining key trajectories of changes in the proteome during normal and disease states. Cluster 1 represented proteins that exhibit low abundance in the control muscle and high abundance in the mutant muscle at both the pre-symptomatic and symptomatic stages. Clusters 2 and 4 represented proteins with high levels in the juvenile stage but a large reduction at the adult stage in the normal muscle. Interestingly, these clusters in the mutant fish showed reduction at both juvenile (pre-symptomatic) and adult (symptomatic) stages. Finally, cluster 3 exhibits protein levels that are elevated in control muscle at both juvenile and adult stages but reduced in mutant muscle at both stages. These data show extensive and dynamic remodeling of the cellular proteome during normal skeletal muscle development. Mutant muscle shows major changes in the proteome compared to the control muscle. Moreover, most of the differential proteomic changes in the mutant muscle compared to the control muscle are observed during the juvenile (pre-symptomatic) state and these changes are mostly static during disease onset and progression. Taken together, this indicates that gene expression is subject to complex post-translational regulation *in vivo* resulting in dynamic remodeling in normal skeletal muscle development. Furthermore, KLHL40 deficiency alters the proteomic trajectories of skeletal muscle during disease onset and progression.

### Bioenergetic and biosynthetic metabolic changes in proteomics precede structural changes in skeletal muscle in KLHL40 deficiency

NM and most of the other skeletal muscle diseases are associated with mitochondrial abnormalities and impairment of cellular metabolic processes during the late pathological stages. To investigate biological pathway associated with KLHL40 deficiency, pathway enrichment analysis was performed on proteins significantly increasing or decreasing in the mutant at each stage (FDR P<0.05; Figure 4A and Figure S4). Proteins showing increased abundance in mutant at both juvenile (pre-symptomatic) and adult (symptomatic) stages in comparison to controls (Cluster 1) were enriched in processes associated with cellular metabolism and developmental muscle proteins. Categories associated with cellular metabolism showed enrichment of amino acid and lipid metabolism, glycolysis, mitochondrial respiration, nucleotide metabolism in mutant muscle at both stages (Figure 4A red and yellow nodes, Table S4). Anaerobic glycolysis and mitochondrial respiration (oxidative phosphorylation, OXPHOS) are the primary regulators of cellular bioenergetics in differentiated skeletal muscle. Glucose uptake (hexokinase) and glycolytic pathway enzymes (Pygma, Pygmb, Eno3, Pfkm, Aldoa, Aldob, Aldoc, Pgk1, Pgam1, Pgam2 and Pkmb) were increased in the mutant muscle during the juvenile stage compared to controls (pre-symptomatic stage). Lactate dehydrogenase, that catalyzes pyruvate to lactate conversion, and thereby slow TCA cycle flux, was also increased in the mutant muscle. TCA cycle enzymes also showed upregulation in the mutant muscle. This suggests that pathways regulating bioenergetics balance in skeletal muscle are upregulated in the mutant muscle. However, downstream enzymes in the electron transport chain were not altered in Klhl40 deficient muscle compared to control muscle (Tables 1 and S4). Mutant muscle also exhibited increased enzyme levels that participate in biosynthetic pathways regulating fatty acid oxidation, amino acid metabolism, nucleotide metabolism and glycogen degradation at the juvenile state (yellow nodes, Figure S4, Table S4). This metabolic shift is similar to increased glycolytic and biosynthetic pathways known as the Warburg effect which is commonly used by proliferative cells and also to maintain the progenitor states (27). Transition to a more differentiated state is associated with an increase in oxidative phosphorylation (28) (29). In parallel, mutant muscle also exhibited upregulation of developmental pathways that are downregulated at both stages in control muscle (Klhl41, Flnc, Unc45, Tnnt2, Mybpc1). These proteins are normally expressed during early sarcomere assembly during primary myogenesis in the fetal stage and indicate either a defect in downstream processing of these proteins or delayed kinetics of sarcomere assembly and differentiation. This suggests that altered bioenergetic and biosynthetic pathways imbalance of the muscle associated with enrichment of early development program likely contribute to the first stage of disease pathogenesis even before structural and functional abnormalities are observed in skeletal muscle.

**Figure 4.**
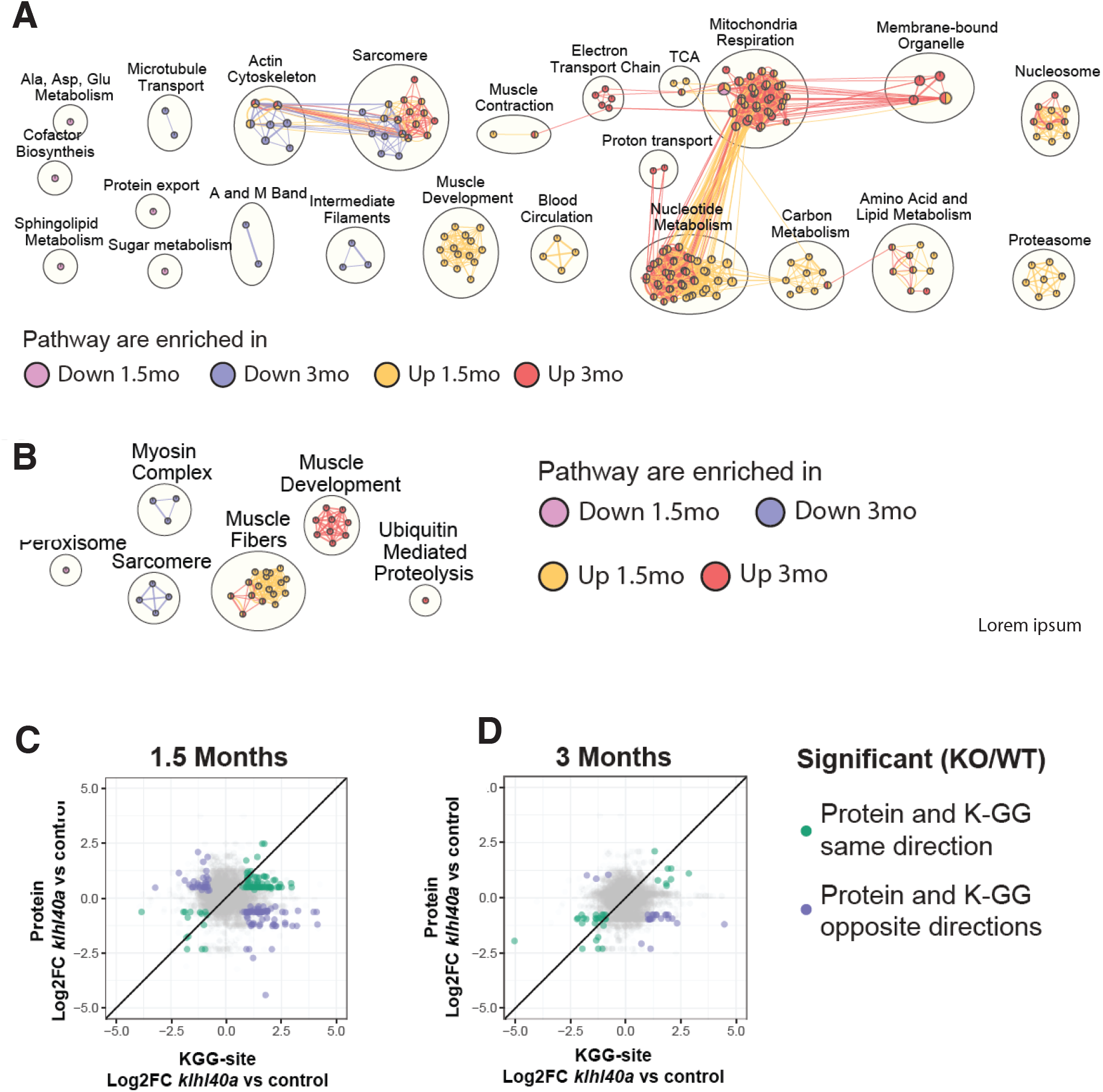
Pathways regulated by changes in proteome and ubiquitylome mediated by KLHL40. (A) Network visualization of pathway enrichment results from *klhl40a KO* differential proteins compared to controls. Nodes (circles) indicate pathways significantly enriched in proteins that increase at 1.5 months, decrease at 1.5 months, increase at 3 months, or decrease at 3 months. Edges (connections) show nodes with overlapping genes. Clusters of nodes summarize pathways with similar biological functions. (B) Network visualization of pathway enrichment results from *klhl40a KO* differential KGG-sites compared to control. Nodes (circles) indicate pathways significantly enriched in KGG-sites that increase at 1.5 months, decrease at 1.5 months, increase at 3 months, or decrease at 3 months. Edges (connections) indicates nodes with overlapping genes. Clusters of nodes summarize pathways with similar biological functions. (C) Fold-changes of KGG-sites and their cognate protein in response to *klhl40a KO* in 1.5 months animals compared to controls. Data points are colored if both the KGG-site and the cognate protein show differential abundance in KO vs (+/+) control and have the same direction (green) or opposite directions (purple). (D) Fold-changes of KGG-sites and their cognate protein in response to *klhl40a KO* in 3 months animals compared to controls. Data points are colored if both the KGG-site and the cognate protein show differential abundance in *KO* vs (+/+) control and have the same direction (green) or opposite directions (purple).

### Reduced Vesicle trafficking and sarcomeric proteins are associated with *klhl40a* mutant muscle

In normal skeletal muscle, clusters 2 and 4 correspond to proteins with high levels in the juvenile stage but a significant reduction at the adult stage (Fig 3B). Interestingly, these clusters in mutants showed reduction at both juvenile and adult stages and represent proteins in lipid catabolic process (e.g., Pck1, Pck2), vesicle trafficking (e.g., Sec16b, Srp14, Timm10) and the UDP-N-acetylglucosamine biosynthetic process indicating the involvement of vesicular trafficking pathway. This was also evident from the accumulation of abnormal vesicles in the *klhl40a* mutant muscles (Figure 2).

Finally, cluster 3 exhibits protein levels that are significantly less abundant in mutant muscle at both stages. Pathway enrichment showed that proteins in this cluster include proteins of the actin cytoskeleton (e.g. TMOD, TNNT3, TNNI2, TPMA, ROCK1) sarcomere assembly (e.g., MYOM, ACTN3, CAPZ, SMYD1) intermediate filaments (e.g. PLEC, DSP, MTM1) and microtubule transport (e.g. KIF5B, MFN2, DYNC2H1) (Figure 3F, blue and cyan nodes). These proteins are required for the formation and maintenance of mature sarcomeres and downregulation in both stages in mutant muscle may contribute to a defect in sarcomere growth in KLHL40 deficiency as observed by electron microscopy (Figure 2). Correlation of protein response to *klhl40a* mutants across juvenile and adult animals showed the proteome is dynamically remodeled during disease progression (Figure 3D, Tables S6-S7). Most of the proteins that are altered at both juvenile and adult stages in the mutant muscle exhibited differential abundance in the same direction and rarely showed differential abundance in opposite directions between pre-symptomatic and symptomatic states. These studies show that KLHL40 deficiency is associated with a decrease in vesicular trafficking and sarcomeric proteins that are required for the formation and maintenance of mature myofibers.

### Quantitative KEGG regulation of proteome in skeletal muscle is required for vesicle trafficking, glycolysis, and sarcomeric proteins

Deep ubiquitylome profiling illuminated changes in ubiquitylation dynamics during skeletal muscle development and disease onset in Klhl40 deficiency. Similar to the dynamics of changes observed in the proteome, changes in the ubiquitylome of mutant muscle are established before functional and structural changes are observed in skeletal muscle (Table S2). A heat map of Hierarchical cluster analysis of ubiquitin sites (Ub-sites) with differential abundance between wild-type and mutants in the ubiquitylome data showed four different clusters classified into two broad categories (Figure 3C). Clusters 1 and 2 exhibited proteins with decreased Ub-sites in both pre-symptomatic and symptomatic mutant states. We expect that many of these proteins are direct targets of ubiquitylation by the Klhl40-Cul3 complex. Previous studies have shown nebulin is a direct ubiquitination target of Klhl40-Cul3 complex and was identified in cluster 1 validating our hypothesis (24). Like the changes we observed for protein abundance levels, most of the changes in ub-sites are restricted to either pre-symptomatic or symptomatic stages. In addition, some proteins exhibited changes in ub-sites at both pre-symptomatic and symptomatic stages in the same direction (Figure 3C and 3E). We did not identify any protein exhibiting significant enrichment of ub-sites in different directions (Figure 3E), suggesting that ubiquitylation marks by Klhl40-Cul3 complex and potentially other ubiquitylation enzymes are robust and unidirectional in the disease state. Clusters 3 and 4 represented proteins that exhibited an increase in ub-sites in the *klhl40a* mutant muscle in comparison to control. As a large number of ubiquitin ligases and deubiquitylases showed differential expression between control and mutant muscle, these proteins could be direct targets of many of these enzymes. Pathway enrichment analysis showed that the most significantly downregulated ub-sites nodes in the mutant muscle were the peroxisome and sarcomere proteins (Fig 4B, blue and cyan nodes; e.g. Ttn, Myha, Myhb, Tpma, Myhc4, Myom1) while nodes that exhibited upregulated ubiquitylated peptides included proteins in muscle development and muscle fibers formation (Obscn, Tnni1, Tmod4, Tpm2, Myom2, Tnni2, Cfl2, Ldb3, Des) and ubiquitin mediated proteolysis (Figure 4B, red and yellow nodes). Integration with the proteome data revealed that these highly ubiquitylated proteins have decreased abundance in mutant muscle (Tmod4, Tnni2, Tpm2, Cfl2, Myom2). Many of these highly ubiquitylated proteins are localized to thin filaments and contribute to nemaline myopathy indicating other components of the ubiquitin proteasomal pathways may cause increased ubiquitylation and abnormal degradation of sarcomere proteins in mutant muscle and affect thin filaments stability (Figure 4 C-D, Table S8-S9). Finally, analysis of fold changes of reduced Ub-sites and abundance of their cognate proteins in response to Klhl40 deficiency in opposite directions identified Sar1ab (vesicle trafficking protein), glycolytic proteins (Pkmb, Aldoa, Aldob) and sarcomeric proteins (Ttn, Tnnt2, Nckipsd) with highest differences between the control and mutant muscles suggesting that these could be the direct target of Klhl40-Cul3 mediated ubiquitylation and subsequent protein degradation *in vivo*.

### Dynamic regulation of vesicle trafficking by ubiquitylation regulates skeletal muscle development

The vesicle trafficking pathway is a central pathway in cells for the transport of cargo and secretory proteins. Proteins of anterograde ER-Golgi vesicle trafficking pathways (Sec23a, Sec23b, Sec23d, Golga2) were significantly downregulated in the mutant muscle except Sar1ab (secretion associated Ras GTPase, orthologue of human SAR1A) that showed significantly increased level in the mutant muscle by proteome analysis (Figure 5A). Sar1ab also showed decreased ubiquitylation in the mutant muscle and therefore, could be a direct substrate of the Klhl40-Cul3 ubiquitylation complex (Table S2). SAR1A is a small GTPase that is required for assembly of COPII vesicles at endoplasmic reticulum exit sites (ERES) by recruiting the SEC23/24 complex (30). SAR1A regulates the packaging of specific cargo proteins into vesicles at the ER for export to the Golgi apparatus by shuttling from the cytoplasm to the ER during protein trafficking. SAR1A is expressed at low levels in normal skeletal muscle and previous *ex vivo* studies have shown that overexpression of SAR1 results in excessive membrane tubulation (31). However, the physiological roles and effects of SAR1A perturbations in skeletal muscle are not known. Electron microscopic examination of skeletal muscle in *klhl40a* mutant muscle (3 months) revealed abnormal, excessive membrane tubulation in the SR-ER region that was absent in the control skeletal muscle suggesting that Sar1a upregulation in the mutant muscle could be a key to influencing muscle pathology (Figure 2A-B). To investigate the possible role of Sar1a upregulation on the disease pathology in Klhl40 deficiency in skeletal muscle, western blot analysis was performed to validate the findings of the proteomics data. Western blot analysis in control and mutant *klhl40a* mutant skeletal muscle protein extracts at adult stage (3 months) confirmed increased Sar1a protein levels in the mutant skeletal muscle in comparison to the WT control (Figure 5B). In normal conditions, Sar1a recruits the Sec24d protein complex at ER exit sites to generate functional COPII-coated vesicles for cargo delivery to the Golgi from the ER. Western blot analysis validated that in mutant muscle, Sec24d protein levels were also significantly reduced in comparison to control muscle suggesting that excess tubulation in mutant muscle does not correspond to the assembly of functional COP-II vesicles (Table S1, Figure 5B-C). As COP-II vesicles are transported to the Golgi apparatus for cargo sorting and extracellular secretion, we observed that key Golgi protein, Golga2, was significantly reduced in the mutant muscle in comparison to control muscle further validating Golgi defects in the mutant muscle that was further confirmed by Western blot. Finally, to test if SAR1A upregulation in skeletal muscle is the direct cause of vesicle trafficking defects observed in Klhl40 deficiency, human *SAR1A* mRNA was overexpressed in the wild-type zebrafish. Ultrastructural examination of zebrafish larvae by electron microscopy showed that *SAR1A* mRNA overexpression (50-100ng) resulted in abnormal membrane tubulation in the SR-ER region similar to *klhl40a* mutant fish (Figure 5D). In addition to membrane tubulation defects, high concentration of SAR1A (100ng) resulted in thickening of Z-lines (Figure 5D, arow), a pathology associated with nemaline myopathy.

**Figure 5.**
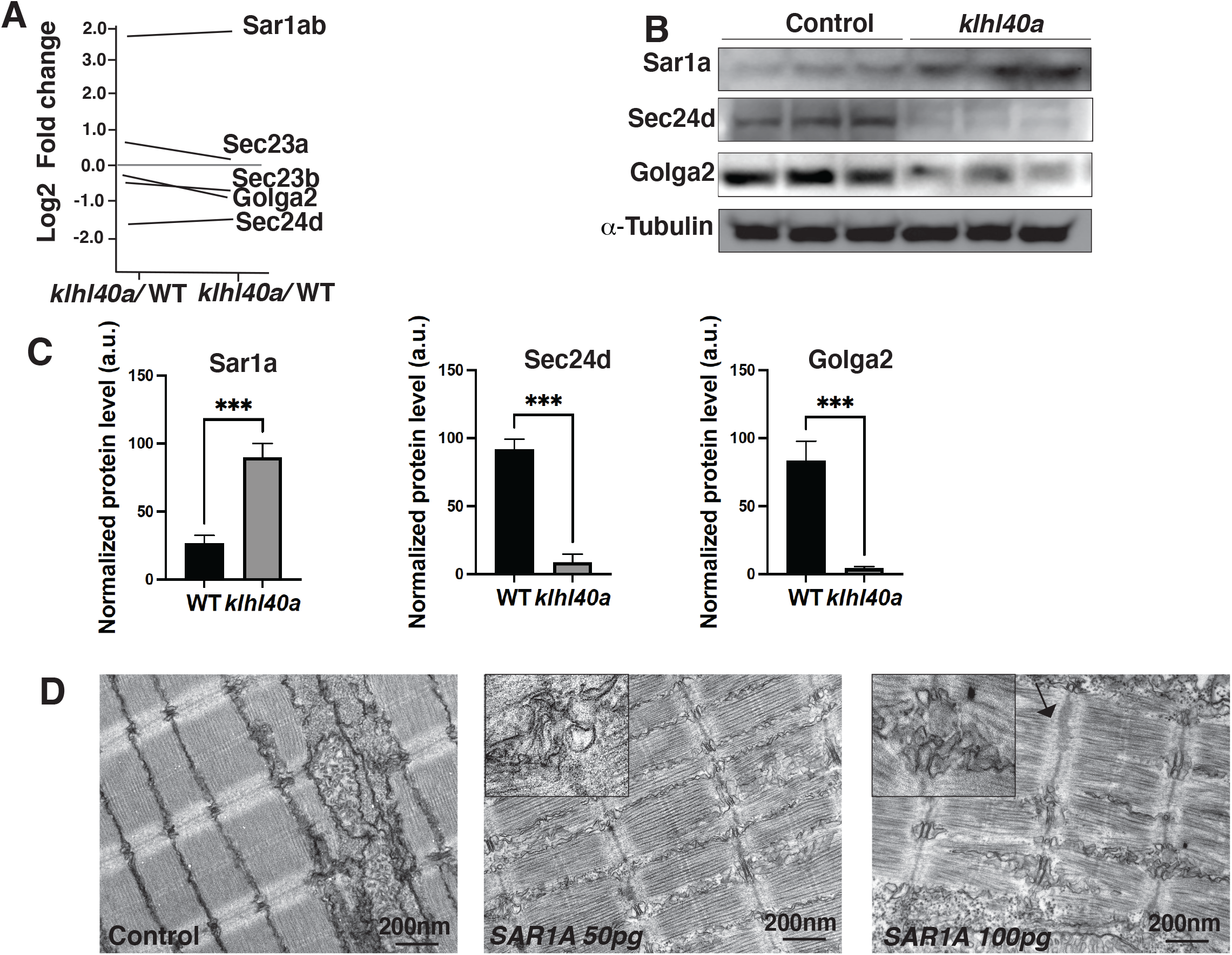
KLHL40 loss result in perturbation of ER-Golgi vesicle trafficking pathway through secretion associated Ras GTPase (Sar1a). (A) ER-Golgi vesicle trafficking proteins exhibit altered levels in *klhl40a* mutant muscle as compared to control (+/+) in proteome analysis. (B) Western blot showing ER-exit site protein Sar1a is upregulated in *klhl40a* mutant muscle and downstream COPII and Golgi proteins are downregulated in mutant muscle (3mo) (C) Quantification of the protein by Western blot in *klhl40a* and control zebrafish. N=3 in each group. Data are mean ± s.e.m; with one way analysis of variance (ANOVA) and Tukey’s HSD test (****p<0.001). (D) Transmission electron microscopy (TEM) of zebrafish larval fish (4 dpf) with *SAR1A* mRNA overexpression demonstrating abnormal vesicle formation in the SR-ER region.

To examine if the increased amount of Sar1a protein is associated with changes in the protein localization in mutant skeletal muscle, immunofluorescence analysis was performed in the control and *klhl40a* skeletal muscle. Immunofluorescence showed that Sar1a protein exhibited ubiquitous localization mostly in the sarcoplasm of the control muscle. In comparison, Sar1a is redistributed in a striated pattern in the mutant skeletal muscle in proximity with the ER-SR marker protein (Ryr1) (Figure 6). This suggest in Klhl40 deficiency, Sar1a is localized to SR-ER region. These studies provide evidence that abnormally increased amount of Sar1a-mediated changes contribute to abnormal vesicle formation, changes in key protein regulators of ER-Golgi vesicle trafficking and contributes to disease pathology.

**Figure 6.**
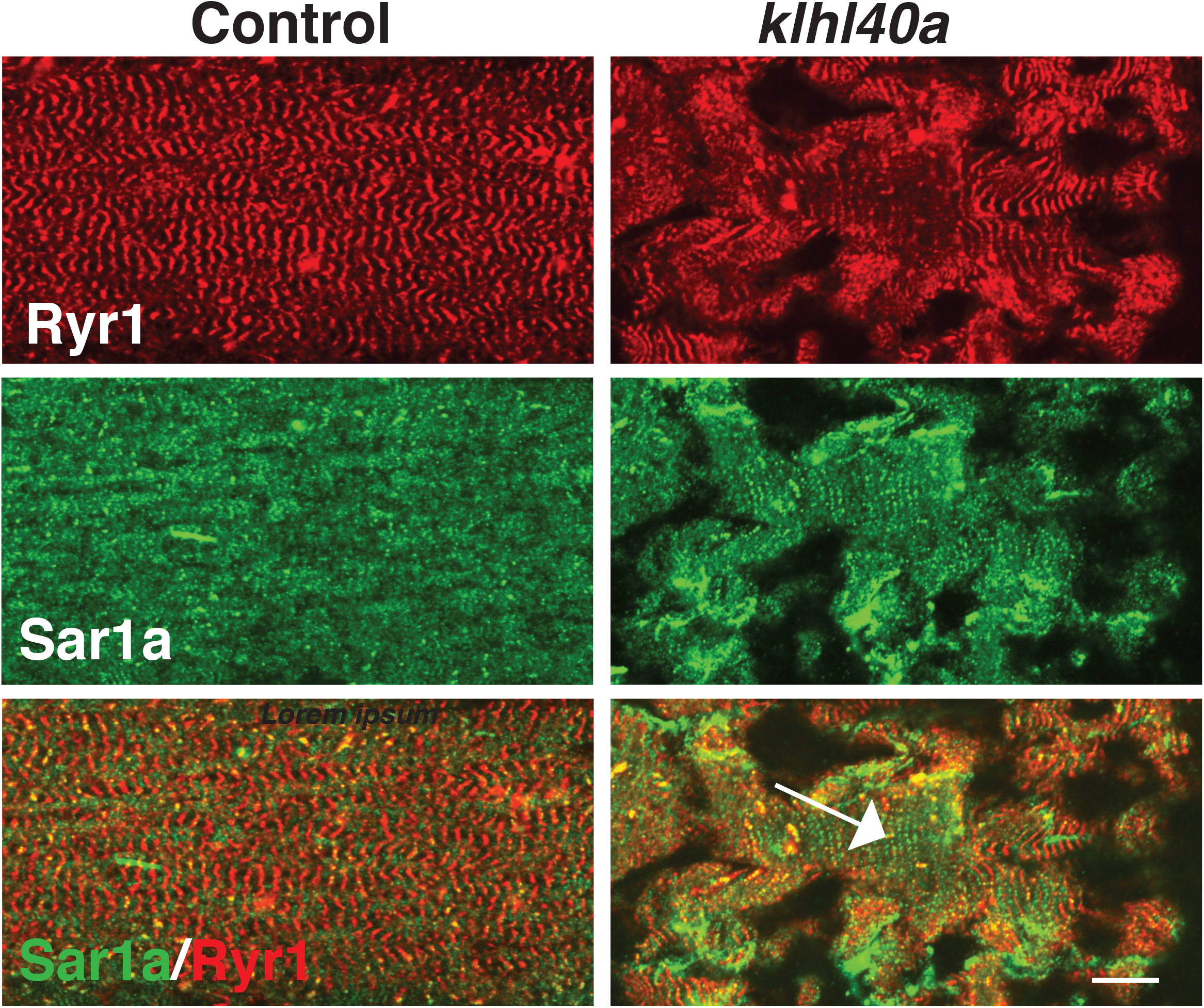
Abnormal localization of sar1a to SR-ER region in KLHL40 deficiency. sar1a is localized to sarcoplasm in the muscle fibers in control muscle and co-localized with ryr1 in *klhl40a KO* muscle (arrow).

### KLHL40-CUL3 regulates SAR1A levels through ubiquitylation

KLHL40 is a substrate-specific protein for CUL3 E3 ubiquitin ligase that targets specific protein substrates for ubiquitination which affects the stability of the target proteins. To test if SAR1A is a direct substrate of KLHL40, we performed co-immunoprecipitation assays in C2C12 cells that showed SAR1A is a direct interactor of KLHL40 (Figure 7A). As SAR1A protein is increased in KLHL40 deficiency, we evaluated if there is a direct reciprocal interaction between KLHL40 and SAR1A proteins by overexpression assays in C2C12 cells. A gradual decrease in KLHL40 protein levels resulted in concomitant increase in SAR1A protein levels in muscle cells (Figure 7B). To understand if the interaction between KLHL40 and SAR1A results in reduced levels of SAR1A through ubiquitylation mediated protein degradation, we evaluated the effect of KLHL40 protein on SAR1A stability in the presence of the proteasome inhibitor MG132 (Figure 7C). In the presence of MG132, increased stability of SAR1A was observed suggesting that KLHL40 targets SAR1A for degradation through proteasomes. Analysis of vertebrate SAR1A protein sequences, revealed that SAR1A ubiquitylation site identified by ubiquitylome analysis is highly conserved in vertebrates suggesting SAR1A ubiquitylation in skeletal muscle may be conserved in all vertebrates (Figure 7D). Finally, to examine whether the KLHL40-CUL3 complex can directly promote SAR1A ubiquitination, *in vitro* ubiquitination assay was performed with neddylated CUL3 and recombinant SAR1A protein with wild type or disease causing KLHL40 proteins (Figure 7E-6F). Western blot analysis showed an increase in SAR1A ubiquitylation as a function of time in the presence of wild-type KLHL40 (Figure 7F). To understand the role of disease-causing KLHL40 missense variants in disease pathology through SAR1A-mediated pathways, ubiquitylation of SAR1A by NM causing KLHL40 missense variants was also studied. Variants in the N-terminal BTB domain of KLHL40 (L86P) and BACK domain (W201L) showed similar SAR1A ubiquitylation as wild-type protein but variants in the Kelch domains (R311L and E528K) resulted in a significant reduction in SAR1A ubiquitylation (Figure 7F-G). As Kelch proteins bind their targets through the C-terminal Kelch domains, this suggests that patients with loss of function or missense variants in the Kelch domains in KLHL40 may exhibit reduced SAR1A ubiquitylation. Finally, SAR1A ubiquitylation was evaluated in C2C12 cells in the presence or absence of KLHL40 with CUL3 E3 ubiquitin ligase by overexpression and immunoprecipitation assay. This showed that KLHL40 is required for SAR1A ubiquitination in the context of muscle cells by CUL3, as no ubiquitylation was observed in the absence of KLHL40 (Figure 7H). Together, these results demonstrate that KLHL40-CUL3 is a negative regulator of SAR1A in skeletal muscle under normal conditions.

**Figure 7.**
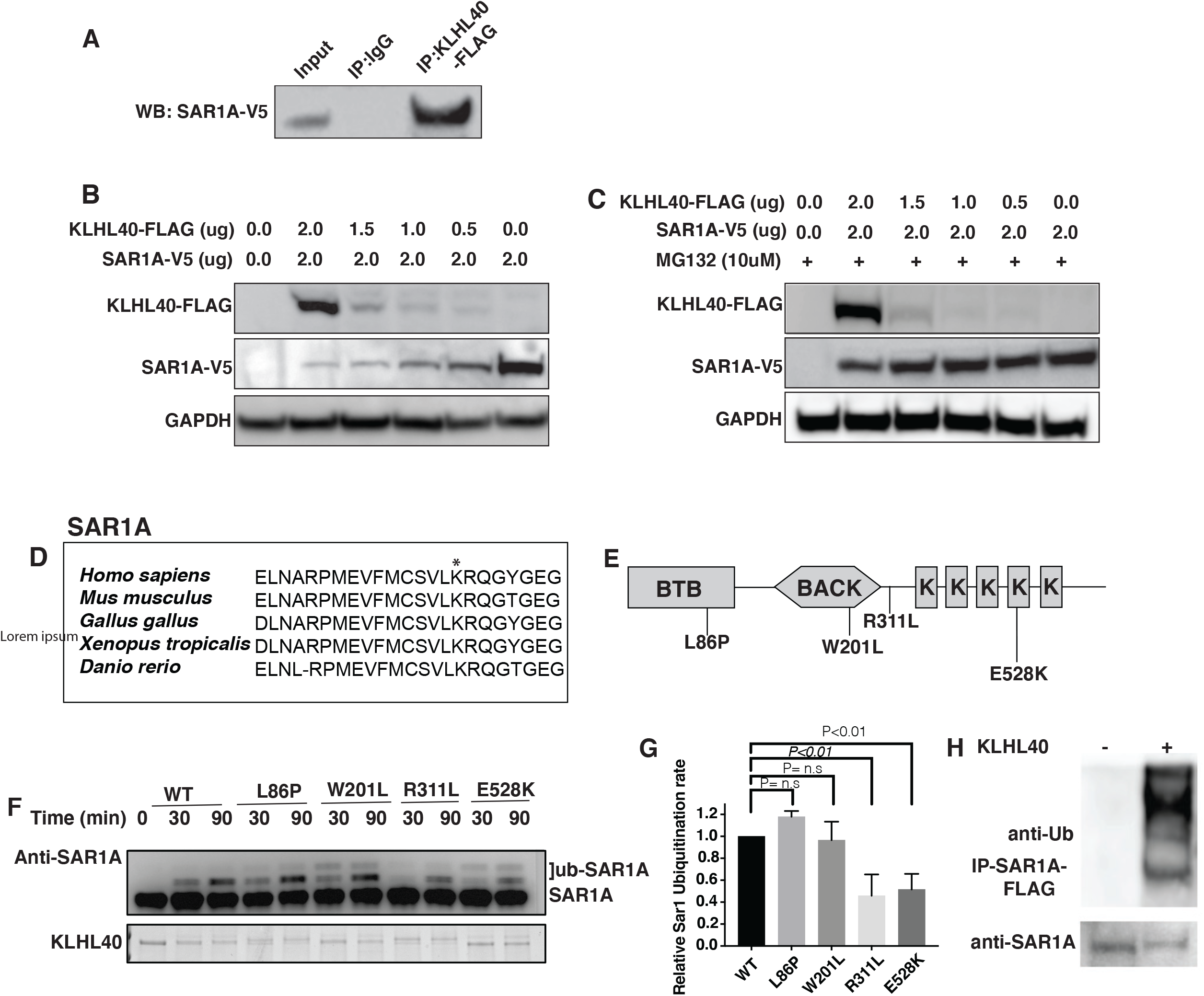
SAR1A is a direct ubiquitylation target of KLHL40-CUL3 complex and differently ubiquitylated by disease causing mutation in *KLHL40*. (A) Coimmunoprecipitation in C2C12 cells showing KLHL40 directly interacts with SAR1A. (B) Co-overexpression of decreasing amounts of KLHL40-FLAG and constant amount of SAR1A-V5 in C2C12 myoblasts demonstrates that KLHL40 is a negative regulator of Sar1A protein. (C) Co-overexpression of decreasing amounts of KLHL40-FLAG and constant amount of SAR1A-V5 in C2C12 myoblasts in the presence of UPS inhibitor MG135 increases the Sar1a protein levels in comparison to MG132-condition. (D) Alignment of amino acid sequence of SAR1Aubiquitylation site demonstrates high conservation in vertebrates. Lysine residue (K) is marked by the asterix. (E) Localization of different disease-causing variants in KLHL40 in the different protein domains. (F) *In vitro* ubiquitylation of human SAR1A by CUL3 protein complex in the presence of wild-type and disease causing KLHL40 proteins. (G) Quantification of the relative human SAR1A ubiquitylation by Wild-type and disease causing KLHL40-CUL3 complex. (H) Ubiquitylation of overexpressed SAR1A in the presence of KLHL40 in C2C12 myoblasts.

### Trafficking of extracellular proteins is perturbed in *klhl40a* mutant muscle

Defects in ER-Golgi trafficking underlie many skeletal muscle diseases, but many proteins that are trafficked by the ER-Golgi cargo vesicles still remain to be identified in skeletal muscle. COP-II vesicles are essential for the transport of large cargo proteins such as procollagens. KLHL40 deficient skeletal muscle exhibited abnormalities in the ECM region in mutant fish (Figure 2). To understand if the abnormal ECM in KLHL40 deficiency is caused by defects in procollagen trafficking to ECM, immunofluorescence was performed on frozen skeletal muscle sections of *klhl40a* mutant and control zebrafish with a procollagen antibody. Immunofluorescence analysis showed extensive immunoreactivity of procollagens intracellularly in *klhl40a* mutant muscle in comparison to the control muscle (Figure 8). In addition, integrin beta-1, an ECM protein, was significantly reduced in ECM in the *klhl40a* mutant muscle in comparison to the control muscle suggesting defects in ECM in the mutant muscle. This suggests a defect in the trafficking of collagen proteins in KLHL40 deficient muscle may underlie ECM defects.

**Figure 8.**
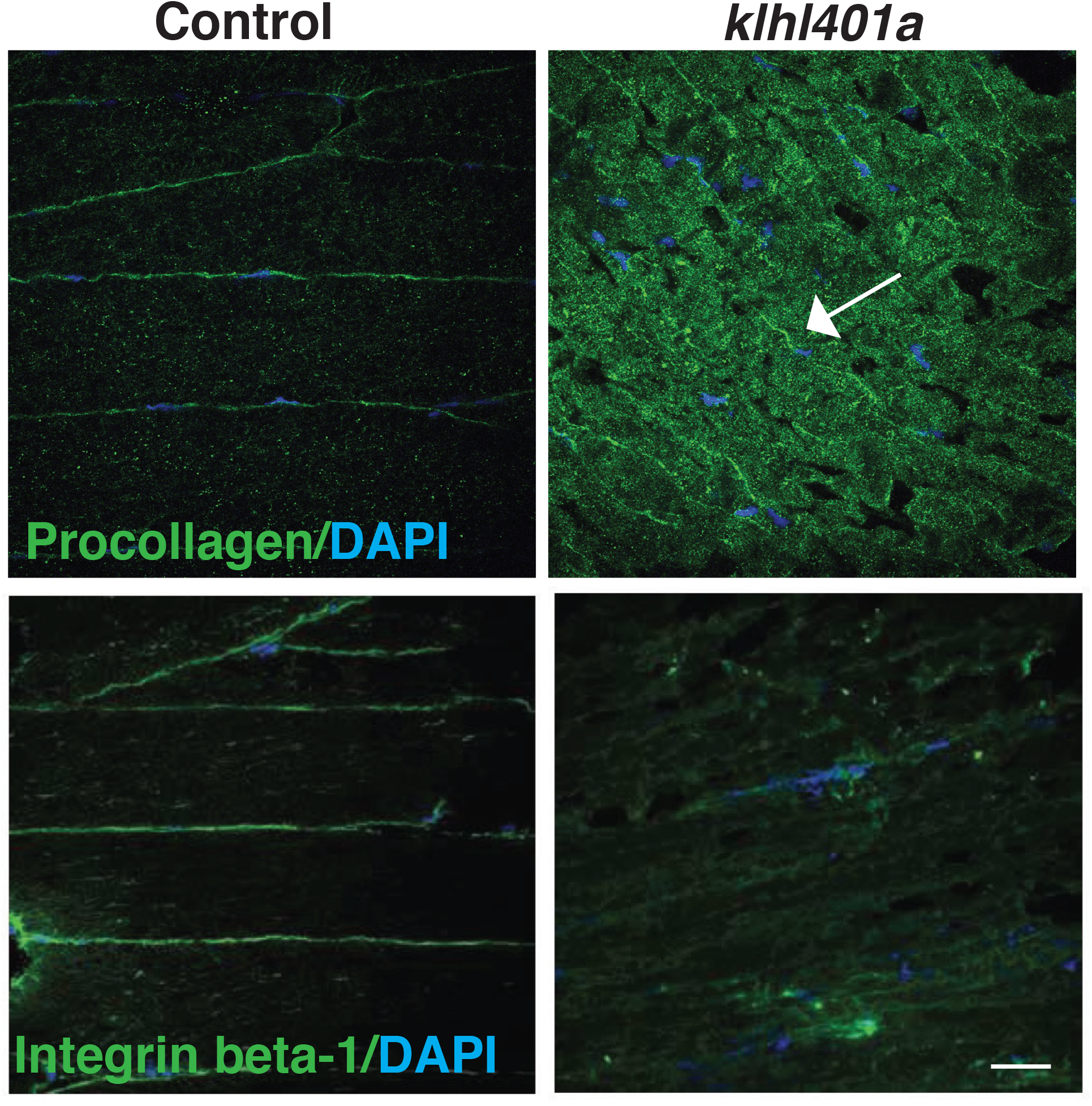
Abnormal trafficking of collagens to extracellular matrix in KLHL40 deficiency. Immunofluorescence of procollagen and integrin in control and *klhl40a KO* muscle. Mutant muscle displays intracellular accumulation of procollagen in *klhl40a KO* muscle in comparison to +/+ control. Integrin level is reduced in the ECM in the mutant muscle.

### KLHL40 human patients exhibit vesicle accumulation and ECM defects

To investigate if SAR1A upregulation and collagen accumulation is associated with disease pathology in KLHL40 deficiency, skeletal muscle of NM patient *KLHL40* patient (c.46C>T, p.Gln16*) was examined with immunofluorescence. Immunofluorescence analysis of skeletal muscle of *KLHL40* NM muscle showed increased immunoreactivity for SAR1A protein in most of the myofibers. Many myofibers exhibited very high level of SAR1A immunoreactivity (Figure 9A, white arrows) compared to control muscle. KLHL40 deficient skeletal muscle also showed increased amount of collagen within the muscle fibers similar to *klhl40a* fish (Figure 9A, arrowheads). Analysis of the ultrastructure of skeletal muscle from KLHL40 patient (c.[932G>T];[1516A>C] p.[Ag311Leu];[Thr506Pro] revealed that in addition to nemaline bodies, extensive vesicle accumulation (arrows) and aberrant ECM structures with reduced collagen fibers (arrowhead) similar to Klhl40 deficient zebrafish were observed (Figure 9B). This suggests that defects in vesicle trafficking contribute to disease onset and pathology in KLHL40 deficiency in patients’ skeletal muscle.

**Figure 9.**
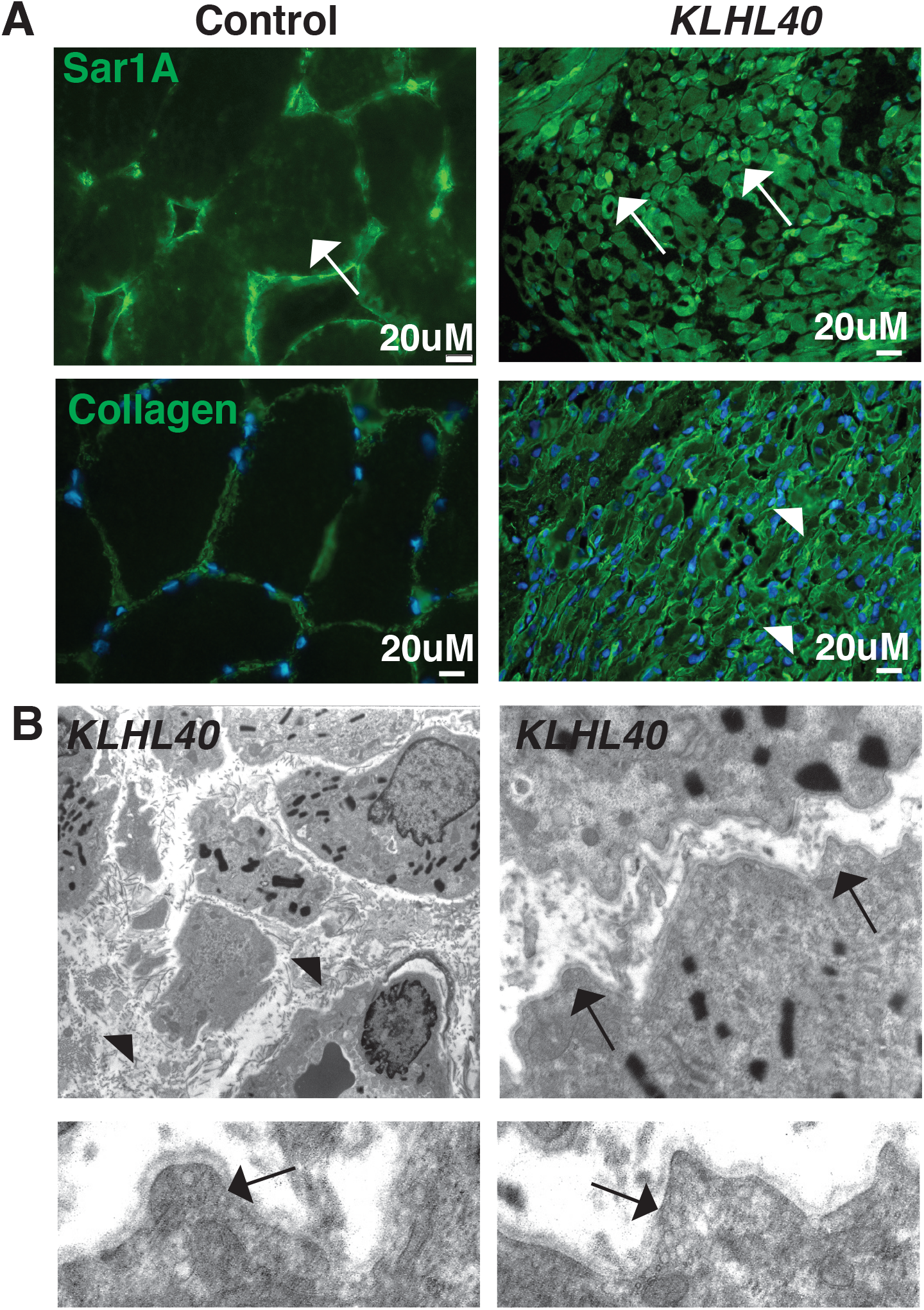
KLHL40-NM patients exhibit increased Sar1a protein and vesicle accumulation with ECM defects in skeletal muscle. (A) Immunofluorescence in control and *KLHL40* patients muscle biopsy showing increased SAR1A protein in patients’ muscle (white arrows). Moreover, increased accumulation of collagen is seen in patient’s muscle (white arrowhead). (B) Transmission electron microscopy of KLHL40 patient muscle showed vesicle accumulation (arrows) and disorganized and damaged extra cellular matrix between myofibers (arrows).

## Discussion

Skeletal muscle development is a highly coordinated process that involves differentiation of muscle stem cells, fusion of myoblasts and formation of multinucleated myotubes and finally, the development of sarcomeres. In parallel, an extensive intracellular membrane network is established in juxtaposition to sarcomeres to produce force-generating myofibers. Rapid fine-tuning of cellular phenotypes to support the dynamic transitions is accomplished through post-transcriptional and post-translational processes. These control rates of protein synthesis and modify protein functions and dynamic degradation of the proteins through the UPS. The protein degradation process during skeletal muscle growth and disease onset is highly selective as evident from the identification of mutations in components in the UPS pathway in human myopathies that perturb specific stages of muscle development and growth and result in impaired motor function (15, 32, 33). In particular, variants in *KLHL40* and *KLHL41* result in severe forms of congenital myopathy with neonatal lethality with extensive sarcomeric disarray and contractures (3, 13). However, *in vivo* studies into proteome remodeling during skeletal muscle development and how disease-causing genes in the ubiquitylation pathway alter cellular pathologies are lacking.

To address these gaps and identify *in vivo* events that initiate the pathology and the mechanisms through which these events become pervasive in KLHL40 deficiency, we performed quantitative global proteome and ubiquitylome analysis in skeletal muscle from non-symptomatic stages to symptomatic stages of disease progression (25, 26, 34). We identified that skeletal muscle proteome under the normal state, exhibited plasticity and was dynamically remodeled during growth. In contrast, KLHL40 deficient skeletal muscle exhibited a highly altered proteome during early stages of skeletal muscle growth, (i.e., during pre-symptomatic stages) that remained mostly static during the transition from juvenile to adult stages and throughout disease progression. In addition, the proteome data identified early preclinical signatures suggesting disease-causing processes are established during pre-symptomatic stages before structural and functional deficits are observed in skeletal muscle. Moreover, most of the proteome (78%) that exhibited changes during the juvenile to adult transition did not show significant changes in gene expression by RNA-sequencing. Transcript abundance is assumed to be the main determinant of protein abundances. However, recent studies demonstrate that regulatory processes such as post-transcription, post-translation and protein degradation that impact steady-state protein abundance after mRNA is made (35). Protein abundance regulation also mirrors specific biological roles as regulatory proteins are produced and degraded very rapidly in response to a stimulus, whereas structural or house-keeping proteins are much longer lived (36). Our studies identified extensive changes in bioenergetic and biosynthetic pathways, downregulation of vesicle trafficking and sarcomeric proteins. These changes are established during pre-symptomatic stages of disease development before the emergence of structural defects thus identifying molecular processes that are under extensive post-translational control. Changes in these molecular processes during pre-symptomatic stages are associated with corresponding structural defects in skeletal muscle during symptomatic stages in mitochondria, membrane vesicles and sarcomere size, thus contributing to disease onset and pathology. In addition, the ubiquitylome, like the overall protein abundance levels, showed dynamic changes between juvenile and adult states in normal muscle and exhibited distinct and static ubiquitylome of the mutant muscle compared to control muscle. Combined analysis of ubiquitylome and proteome identified potential KLHL40 target proteins in glycolytic pathway, sarcomere development and vesicular trafficking.

Skeletal muscle is dependent on anaerobic glycolysis and oxidative phosphorylation for its bioenergetic demands (37). During skeletal muscle development, aerobic glycolysis is the primary source of bioenergetics in the proliferative cells, but this changes to OXPHOS during differentiation. Our combined analysis of the ubiquitylome and proteome identified many glycolytic enzymes that exhibited reduced ubiquitylation and increased protein levels in KLHL40 deficiency. This suggests that KLHL40 directly or indirectly acts as a negative regulator of glycolysis in skeletal muscle and contributes to a perturbed bioenergetic state in the mutant muscle. Pyruvate kinase M2 (Pkm2) High levels of glycolytic enzyme pyruvate kinase (Pkm2) contributes to skeletal muscle atrophy in myotonic dystrophy (38). While ultrastructure evaluation did not reveal atrophied muscle in *klhl40a* zebrafish mutants, we identified key muscular atrophy factor MuRF1 (Trim63) was significantly upregulated during pre-symptomatic and symptomatic stages in the mutant muscle. This suggests that perturbation of the glycolytic pathway may eventually contribute to muscle damage in KLHL40 deficiency. Future studies may provide better mechanistic insights into upstream signals that direct the ubiquitylation of glycolytic enzymes and modulate their stability by KLHL40 during normal and disease states.

Our studies identified that KLHL40 regulates skeletal muscle growth and function through negative regulation of vesicle trafficking by ubiquitylation. Vesicle trafficking is coordinated by multiple organelles that coordinate transport of proteins synthesized in the ER to the extracellular space or other cellular compartments through the Golgi apparatus (39). Currently, a detailed understanding of vesicle trafficking in skeletal muscle is lacking. Human genetic studies have identified pathogenic variants in key proteins in the vesicular trafficking pathway that result in myopathies in affected patients. Variants in *GOLGA2*, a Golgi protein, are associated with neurodevelopmental abnormalities associated with myopathy (40, 41). Autosomal dominant mutations in *BICD2* cause spinal muscular atrophy lower extremity predominant 2 and/or myopathy (4, 42, 43). BICD2 interacts with Rab GTPase (Rab6) to regulate the trafficking of constitutive cargos from the trans-Golgi network to the plasma membrane (44). Recent work has identified mutation in *BET1* as a cause of congenital muscular dystrophy associated with epilepsy. BET1 is required for fusion of ER-derived vesicles with the ER-GOLGI intermediate compartment and suggested to perturb the ER-Golgi trafficking in affected patients (45). Variants in a number of genes associated with congenital muscular dystrophies (*POMT1, POMT2, TRAPPC11, GOSR2*) encode proteins that are localized to different membrane compartments of the vesicle trafficking pathway (46-49). The potential of improving vesicle trafficking to rescue disease pathology skeletal muscle disorders has also been addressed. Overexpression of thrombospondin 4 (THBS4) improves muscle function in muscular dystrophy by improving vesicle trafficking of dystrophin-glycoprotein and integrin attachment complexes to stabilize sarcolemma (50). Despite this detailed information on genetics and therapeutic contributions, detailed mechanisms of the vesicular trafficking pathway in skeletal muscle are lacking. How defects in specific components in the trafficking pathway affect the distribution of secretory and extracellular proteins at a local or global scale is not yet well-understood. Nevertheless, our studies provide *in vivo* insights into the requirement of vesicle trafficking in the transport of ECM proteins to maintain healthy skeletal muscle. KLHL40 acts as a negative regulator of this process through ubiquitylation mediated protein degradation of Sar1a, which is required for budding of COPII vesicles from ER to transport large ECM proteins. Our studies further show that defects in COPII and Golgi vesicles associated with Sar1a alterations related to KLHL40 mutations has functional implications for inter-organelle communication. This is evident by a defect in procollagen trafficking to extracellular matrix in KLHL40 deficiency resulting in structural defects in the ECM. A key clinical feature of KLHL40 NM is the presence of contractures in most patients. Defects in ECM are directly associated with contracture in many forms of neuromuscular diseases. Vesicle trafficking dysregulation is associated with neurodegenerative and skeletal disorders and our findings open new avenues on this pathway in neuromuscular diseases.

Nemaline myopathy is associated with defects in sarcomere structure, and it is not clear if formation or growth of sarcomere is affected in NM patients. Our studies demonstrate that in KLHL40-NM, sarcomeres are initially formed normally, but further sarcomere growth is perturbed and provides mechanistic insights into disease pathology. Previous *in vitro* studies have demonstrated that KLHL40 mediated ubiquitylation of sarcomeric protein NEB and LMOD3 improved protein stability in normal muscle (24). We identified multiple ub-sites for NEB in zebrafish skeletal muscle, some of these ub peptides were upregulated for several lysine residues in nebulin and downregulated for other lysine residues in the mutant muscle. However, no changes were seen in the protein stability and ubiquitylation for LMOD3 in *klhl40a* mutant muscle. The reduced amounts of LMOD3 observed in previous study could be due to protein analysis at the terminal stages of disease pathology. Finally, we did not identify proteins that showed both reduced stability and reduced ubiquitylation in KLHL40 deficiency. This suggests that KLHL40 functions to regulate turnover of cellular proteins through ubiquitylation mediated protein degradation. Moreover, multiple ubiquitylation sites on nebulin indicate the involvement of other ubiquitin ligases in regulating the stability and function of muscle proteins and demonstrates complexity of the UPS pathways in skeletal muscle.

Overall, we provide a comprehensive temporal proteomic landscape during skeletal muscle growth and disease development and dynamic fine-tuning of the cellular proteome by ubiquitylation is critical for muscle function. A better understanding of the natural history of the preclinical stage, the evolution of pathophysiology, and structural changes in the muscle during disease progression is needed to develop effective therapies. Our approach resulted in the identification of previously unknown disease pathways. Given the several pathways in which KLHL40 is involved, approaches aiming to inactivate pro-disease pathways and activate protective pathways may be a promising therapeutic strategy for at least this form of NM.

## Methods

Please see the Supplemental methods for a detailed description of generation of zebrafish lines, methodology for proteome and ubiquitylome quantification and pathway analysis, antibodies, plasmids, cell culture, ubiquitylation assays and statistical analyses.

## Acknowledgements

We would like to thank Dr. Nigel Laing for critically reading this manuscript and valuable suggestions. This work was supported by NIH R56AR077017 (VAG), R37GM62437 (PAC), A Foundation Building Strength grant and Innovation Evergreen Fund Award (VAG) and NIH F32HL154711 (PMJB). GR is supported by an Australian NHMRC EL2 Investigator Grant (APP2007769). This work is also supported by an NHMRC Ideas Grant to GR and NL (APP2002640). The DSHB antibody (8c8) developed by (Hausen and Gawantka) was obtained from the Developmental Studies Hybridoma Bank, created by the NICHD of the NIH and maintained at the University of Iowa, Department of Biology, Iowa City, IA 52241.

