## Supplemental Data for "Dynamic regulation of inter-organelle communication by ubiquitylation controls skeletal muscle development and disease onset"

Figure S1

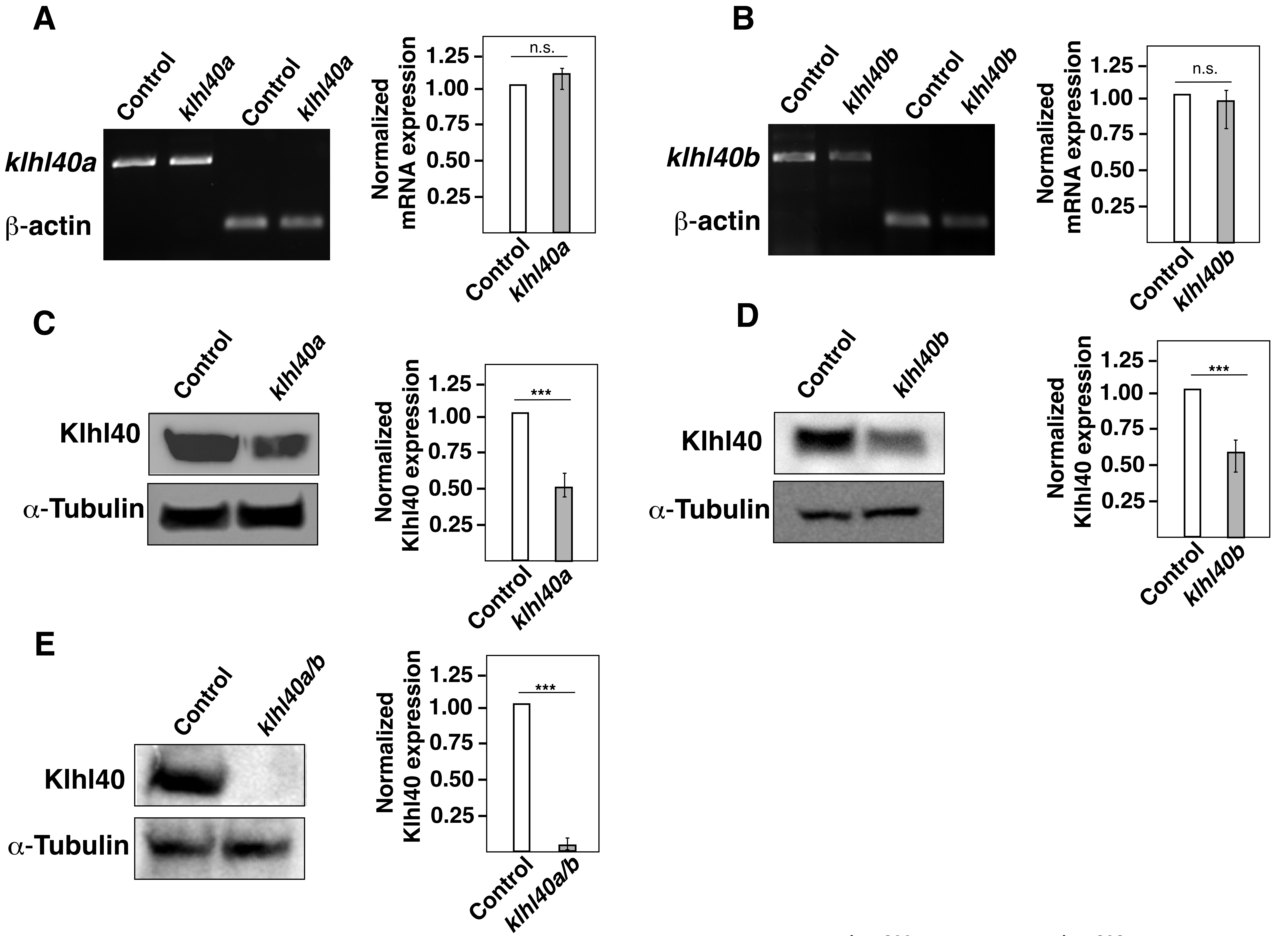


**Figure S1. Quantification of KLHL40 mRNA and protein in *klhl40a$^{bwg200}$* and *klhl40b$^{bwg202}$* alleles (3 months).** (A) RT-PCR analysis of control and *klhl40a$^{bwg200}$* skeletal muscle (B) RT-PCR analysis of control and *klhl40b$^{bwg202}$* skeletal muscle. (C) Western blot analysis of protein extracts from control and *klhl40a$^{bwg200}$* skeletal muscle (D) Western blot analysis of protein extracts from control and *klhl40b$^{bwg202}$* skeletal muscle (E) Western blot analysis of protein extracts from skeletal muscle of control and *klhl40a$^{bwg200}$/klhl40b$^{bwg202}$* double mutants.

### Figure S2

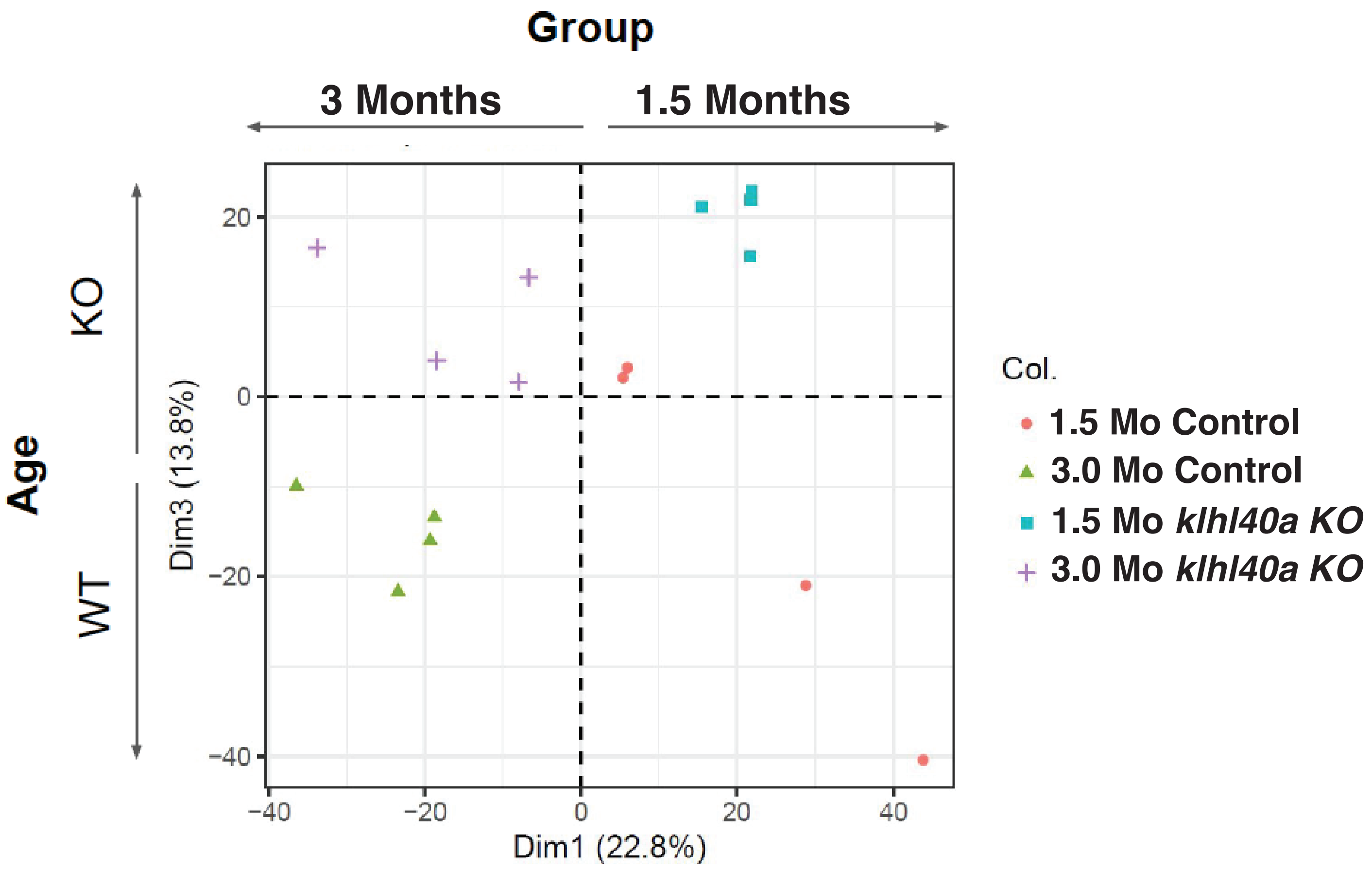


**Figure S2. PCA plot of proteomics data shows clustering of normal and disease groups across ages**

### Figure S3

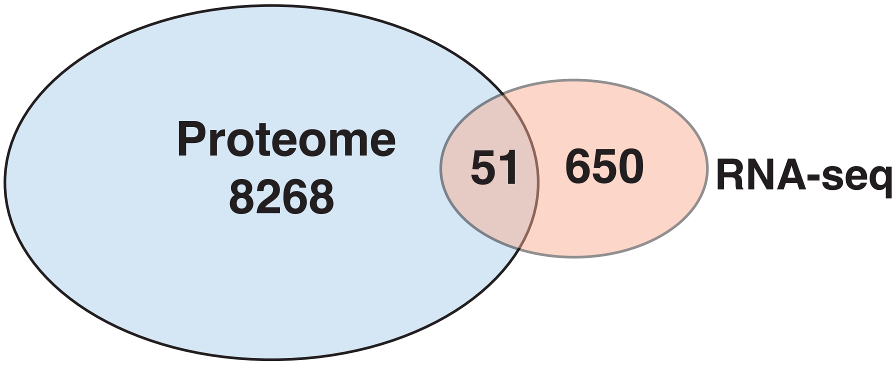


**Figure S3. Overlap of proteomics and transcriptome data.** Differential proteins identified between control and *klhl40a*$^{bwg200}$ mutants at 3 months were compared with differential RNA-seq data from control *klhl40a*$^{bwg200}$ mutant skeletal muscle tissue (3 months) by GeneVenn.

#### Figure S4

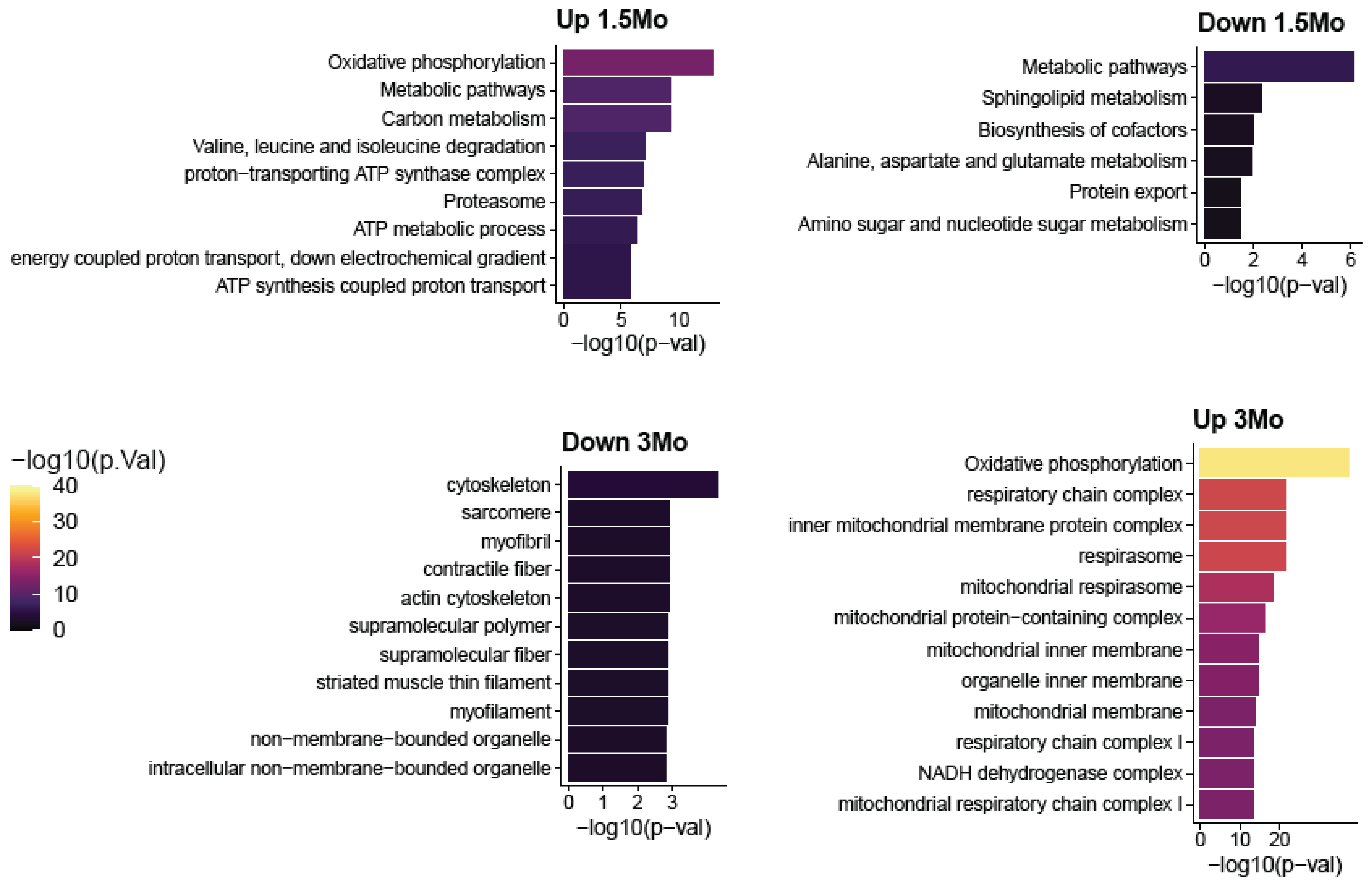


**Figure S4. Top pathway enrichments in proteins that increase or decrease in *klhl40a KO* at 1.5 months and 3 months. Up to 10 pathways per group are shown.**

**Supplemental Table legends**

**Table S1: Differential Proteome expression during disease onset and progression in normal and *klhl40a$^{bwg200}$ KO* at 1.5 months and 3 months.**

**Table S2: Differential Proteome and ubiquitylome changes during disease onset and progression in normal and *klhl40a$^{bwg200}$ KO* at 1.5 months and 3 months.**

**Table S3: Differential transcriptome expression in normal and *klhl40a$^{bwg200}$* at 3 months**

**Table S4: Cluster of enriched pathways by proteome and ubiquitylome analysis at 1.5 months and 3 months.**

**Table S5: Gene and KGG-site clusters changed during transition from 1.5 months to 3 months identified by hierarchal clustering**

**Table S6: Significant changes in proteome in opposite direction at 1.5 months and 3 months**

**Table S7: Significant changes in proteome in the same direction at 1.5 months and 3 months**

**Table S8: Proteome and ubiquitylome correlation in the same direction at 1.5 months**

**Table S9: Proteome and ubiquitylome correlation in different direction at 1.5 months**

**Table S10: Sequences of the sgRNA target sites and primers sequences (5'-3') to clone sgRNAs for creating *klhl40* zebrafish lines.**

#### Supplementary Methods

##### Zebrafish Maintenance and Husbandry

Zebrafish were maintained and bred using standard methods as described (Westerfield, 2000). All experiments and procedures were approved by the Institutional Animal Care and Use Committee at Brigham and Women's Hospital. Wild-type fish were obtained from Tubingen (TU) line and staged by hours (h) or days (d) post fertilization at 28.5°C. Zebrafish embryonic, larval, juvenile and adult stages of development have been described previously (Kimmel, et al., 1995).

##### Creation of zebrafish lines

sgRNAs were designed using the web-based ZiFiT Targeter program (http://zifit.partners.org/) and targeting distinct sites in exon 1 or exon 2 of zebrafish Kelch genes (Sander et al., 2010). The first 2 bp GG sequences at the 5' end of the target sit are a constraint 20 imposed by T7 promoter sequence requirements in addition to the NGG protospacer adjacent motif (PAM) sequence requirement immediately 3' to the target site. Two oligonucleotides, each of 22 nucleotides in length, were used for construction of the guide RNA for each target site. Forward and reverse primers were annealed to create a sgRNA oligonucleotide duplex. Primers sequences are summarized in Table S10. The zebrafish guide RNA expression vector pDR274 was used to create the sgRNAs expression system using the T7 promoter followed by In Vitro Transcription as previously described (Hwang et al., 2013). sgRNAs and Cas9 protein (Thermo Scientific, CA) were co-injected into the yolk sac of one- and two-celled stage zebrafish embryos. Each embryo was injected with a 5ul injection solution containing 2ul of 400ng/nl Cas9 protein and 3ul of 100ng/ul sgRNA. Injected embryos were inspected under the microscope for 3 days and were classified as dead, deformed, or normal phenotypes. Embryos displaying normal phenotype were analyzed to test the efficacy of sgRNAs by identifying target site mutations. To analyze injected fish, genomic DNA was extracted from 6-8 pooled embryos at 3-5 days post fertilization (dpf) and used for DNA sequencing experiments by topo cloning.

##### Identification of Founder Fish and Generation of Isogenic Stable Mutant Fish

Because a fertilized zebrafish embryo develops quickly, direct delivery of sgRNA-Cas9 protein via injection results in chimeric embryos. Founder fish were determined by genotyping tail fin clips of the F0 generation and observing mosaicism at the target site. The mosaic F0 generation were outcrossed to wildtype TU fish for atleast three generations for studies presented in this work. The sequences of sgRNAs are listed in Table S10.

##### In-Solution Digestion

Muscle samples frozen in liquid nitrogen were cryofractured using the cryoPREP tissue disruption system on setting 4 (Covaris). Samples were then lysed for 30 min at 4 °C in urea lysis buffer (8M urea, 50 mM Tris-HCl pH 8.0, 75 mM NaCl, 1 mM EDTA, 2 μg/μl aprotinin (Sigma-Aldrich), 10 μg/μl leupeptin (Roche), and 1 mM phenylmethylsulfonyl fluoride (PMSF) (Sigma-Aldrich)) and cleared by centrifugation at 20,000xg. Protein concentrations were determined by bicinchoninic acid (BCA) protein assay (Pierce), and sample were diluted to a protein concentration of 2ug/ul. Samples were reduced with 5 mM dithiothreitol (DTT) for 1 h at 21°C, followed by alkylation with 10 mM iodoacetamide for 45 min at 21°C. Samples were diluted 50 mM Tris-HCl pH 8.0 to a final urea concentration of 2 M preceding enzymatic digestion. Proteins were digested with endoproteinase LysC (Wako Laboratories) for 2 h at 25 °C followed by overnight digest with sequencing-grade trypsin (Promega) at 25 °C (enzyme-to-substrate ratios of 1:50). Following digestion, samples were acidified to a concentration of 1% formic acid (FA) and cleared by centrifugation at 20,000 rcf. Remaining soluble peptides were desalted using a 100 mg reverse phase tC18 SepPak cartridge (Waters). Cartridges were conditioned with 1 ml 100% acetonitrile (MeCN) and 1 ml 50% MeCN/0.1% FA, then equilibrated with 4X 1 ml 0.1% trifluoroacetic acid (TFA). Samples were loaded onto the cartridge and washed 3X with 1 ml 0.1% TFA and 1X with 1 ml 1% FA, then eluted with 2X 600 µl 50% MeCN/0.1% FA. Peptide concentration of desalted samples was again estimated by BCA assay, dried by vacuum centrifugation.

**TMT labeling of peptides**
TMT labeling was performed as previously described (Zecha et al., 2019). Briefly, a total of 100ug peptides per sample was resuspended in 50 mM HEPES pH 8.5 at a concentration of 5 mg/ml. Dried Tandem Mass Tag (TMT) pro 16-plex reagent (ThermoFisher Scientific) was reconstituted at 20 µg/µl in 100% anhydrous MeCN and added to samples at a 2:1 TMT to peptide mass ratio. The reaction was incubated for 1 hr at 25 °C while shaking and quenched with 5% hydroxylamine to a final concentration of 0.2% for 15 min at 25 °C while shaking. The TMT-labeled samples were then combined, dried to completion by vacuum centrifugation, reconstituted in 1 ml 0.1% FA, and desalted with a 100 mg SepPak cartridge as described above.

**Basic Reverse Phase (bRP) Fractionation**
TMT-labeled peptides were fractionated via offline basic reverse-phase (bRP) chromatography as previously described (Mertins et al., 2018). Chromatography was performed with a Zorbax 300 Extend-C18 column (4.6 x 250 mm, 3.5 µm, Agilent) on an Agilent 1100 high pressure liquid chromatography (HPLC) system. Samples were reconstituted in 900 µl of bRP solvent A (5 mM ammonium formate, pH 10.0 in 2% vol/vol MeCN). Peptides were separated at a flow rate of 1ml/min in a 96 min gradient with the following concentrations of solvent B (5 mM ammonium formate, pH 10.0 in 90% vol/vol MeCN) 16%B at 13min, 40%B at 73min, 44%B at 77min, 60%B at 82min, 60%B at 96min. A total of 96 fractions were collected and concatenated non-sequentially into a final 24 fractions for proteomic analysis. Fractions were dried via vacuum centrifugation and an equivalent of 1ug of peptide was injected for LC-MS/MS analysis.

**Liquid chromatography and mass spectrometry for global proteome analysis**
Dried fractions were reconstituted in 3% MeCN/0.1% FA to an estimated peptide concentration of 1 µg/µl and analyzed via coupled nanoflow liquid chromatography and tandem mass spectrometry (LC-MS/MS) using a Proxeon Easy-nLC 1200 (Thermo Fisher Scientific) coupled to an Orbitrap Exploris 480 Mass Spectrometer (Thermo Fisher Scientific). A sample load of 1 µg for each fraction was separated on a capillary column (360 x 75 µm, 50 °C) containing an integrated emitter tip packed to a length of approximately 25 cm with ReproSil-Pur C18-AQ 1.9 µm beads (Dr. Maisch GmbH).

Chromatography was performed with a 110 min gradient of solvent A (3% MeCN/0.1% FA) and solvent B (90% MeCN/0.1% FA). The gradient profile, described as min:% solvent B, was 0:2, 1:6, 85:30, 94:60, 95:90, 100:90, 101:50, 110:50. Ion acquisition was performed in data-dependent mode with the following relevant parameters: MS1 orbitrap acquisition (60,000 resolution, 350-1800 scan range (m/z), 300% normalized AGC target, 25ms max injection time) and MS2 orbitrap acquisition (20 scans per cycle, 0.7m/z isolation window, 32% HCD collision energy, 45,000 resolution, 50% normalized AGC target, 50 ms max injection time, 15s dynamic exclusion, 50% fit threshold, and 1.2 m/z fit window).

**K-GG enrichment for ubiquitylome analysis**

Ubiquitin enrichment was performed based on the UbiFast protocol (Udeshi et al., 2020). Anti-K-e-GG bead-bound antibodies from the PTM-Scan ubiquitin remnant motif kit (Cell Signaling Technologies #5562) were cross-linked as follows. Beads were washed 3X with 100 mM sodium borate (pH 9.0) and incubated with 20 mM DMP for 30 min at RT. Beads were then washed 2X with 200 mM ethanolamine and incubated overnight at 4°C in 200 mM ethanolamine with end-over-end rotation. Following incubation, beads were washed three times with IAP buffer and stored at 4C at a concentration of 0.5 ug/uL. For each 11-plex experiment, 31.25 ug of cross-linked anti-K-GG bead-bound antibody at 0.5 ug/uL in IAP per channel was aliquoted into 1.5 mL Eppendorf tubes on ice. 1mg peptide per sample was reconstituted to 0.5 mg/mL concentration in IAP buffer and vortexed for 10 min. Peptides were then centrifuged for 5 min at 5,000 g. Each peptide solution was added to a tube of antibody and gently rotated end-over-end at 4°C for 1 h. Following enrichment, samples were centrifuged (1min, 2000 rcf) and the supernatant was removed. Beads were washed with 1.5 mL ice cold IAP followed by 1.5 mL ice cold PBS (30s, 2000 rcf) and reconstituted in 200 uL 100 mM HEPES buffer. For each sample, 400 ug of TMTpro 16-plex labeling reagent in 10 uL acetonitrile was added. Peptides were TMT labeled on-beads while shaking vigorously (1400 rpm) at 20C for 10 min, then quenched with 8 uL 5% hydroxylamine and shaken vigorously for another 5 min washed once with 1.3 mL cold IAP, and again with 1.5 mL cold IAP. Each channel was resuspended and transferred to a combination tube with 130 uL cold IAP. Following combination, each now-empty tube was serially washed with 1.5 mL cold IAP to remove remaining beads, and this 1.5 mL IAP was added to the combination tube and used to wash the combined beads. Combined beads were washed one final time with 1.5 mL ice cold PBS. Once the channels were combined and washed, peptides were eluted twice from the beads by resuspending with 150 uL room temp. 0.15% TFA and incubating 5 min at RT. Each round of acid-eluted K-GG-modified peptides was desalted on an equilibrated two-punch C18 stage tip. Both elutions of K-GG peptides were loaded sequentially, washed 2X with 100 uL 0.1% FA, and eluted into an MS vial with 50 uL 50% ACN/0.1% FA. The eluted peptides were frozen, lyophilized, and reconstituted in 9 uL 3% ACN/0.1% FA, with 4 uL injected twice for two consecutive LC-MS/MS runs.

**Liquid chromatography and mass spectrometry for global proteome analysis**

Reconstituted K-GG enriched peptides were analyzed via coupled nanoflow liquid chromatography and tandem mass spectrometry (LC-MS/MS) using a Proxeon Easy-nLC 1200 (Thermo Fisher Scientific) coupled to an Orbitrap Exploris 480 Mass

Spectrometer (Thermo Fisher Scientific) equipped with a FAIMS interphase. 4 out of 9ul of total eluted material was separated on a capillary column (360 x 75 µm, 50 °C) containing an integrated emitter tip packed to a length of approximately 25 cm with ReproSil-Pur C18-AQ 1.9 µm beads (Dr. Maisch GmbH). Chromatography was performed with a 154 min gradient of solvent A (3% MeCN/0.1% FA) and solvent B (90% MeCN/0.1% FA). The gradient profile, described as min:% solvent B, was 0:2, 2:6, 122:35, 130:60, 133:90, 143:90, 144:50, 154:50. Ion acquisition was performed in data-dependent mode with the following relevant parameters: three FAIMS CV settings (-45V, -50V, and -70V), MS1 orbitrap acquisition (60,000 resolution, 350-1800 scan range (m/z), 100% normalized AGC target, 10ms max injection time) and MS2 orbitrap acquisition (10 scans per cycle, 0.7m/z isolation window, 32% HCD collision energy, 45,000 resolution, 50% normalized AGC target, 120 ms max injection time, 20s dynamic exclusion, 50% fit threshold, and 1.4 m/z fit window).

**Data Analysis**

Raw MS/MS data from heart and liver samples was processed using Spectrum Mill v.7.09.215 (Agilent Technologies). MS2 spectra were extracted from RAW files and merged if originating from the same precursor, or within a retention time window of +/- 60 s and m/z range of +/- 1.4, followed by filtering for precursor mass range of 750-6000 Da and sequence tag length > 0. MS/MS search was performed against the Uniprot *Danio rerio* protein database downloaded on November, 2020 and common contaminants, with digestion enzyme conditions set to “Trypsin allow P”, <5 missed cleavages, fixed modifications (cysteine carbamidomethylation and TMTpro on N-term and lysine), and variable modifications (oxidized methionine, acetylation of the protein N-terminus, pyroglutamic acid on N-term Q, and pyro carbamidomethyl on N-term C). Additional variable modification were added for ubiquitylome (di-glycine residual in K). Matching criteria included a 30% minimum matched peak intensity and a precursor and product mass tolerance of +/- 20 ppm. Peptide-level matches were validated if found to be below the 1.0% false discovery rate (FDR) threshold and within a precursor charge range of 2-6. A second round of validation was then performed for protein-level matches for proteome datasets, requiring a minimum protein score of 13. Ubiquitylome site-centric and protein-centric data, including TMT intensity values and ratio to the median of all samples, was extracted and summarized in a table. Raw mass spectrometry data will be made publicly available in MassIVE upon acceptance of the manuscript.

**Statistical analysis of proteomics data**

Statistical analysis was performed in the R environment for statistical computing. Sample log2 TMT ratios were median centered. Proteins with less than 2 unique peptides were removed from downstream analysis. One sample (4-month knockout replicate 1) was identified as an outlier by PCA and removed from the dataset. In order to identify proteins and KGG-sites with differential abundance between WT and KO groups, a linear model was fit with age and genetic background as experimental factors and moderated T-tests were performed using the limma package (Ritchie et al., 2015). Multiple hypothesis testing correction was performed using the BH method.

**Pathway enrichment and network visualization**
Proteins and KGG-sites showing differential abundance in response to KLHL40 knockout at both ages were used for downstream pathway analysis (adj. p-val < 0.05). Pathway enrichment analysis was performed for features increasing or decreasing in abundance in the KO stain at each of the two ages using the g:profiler tool (Raudvere et al., 2019). The background list of proteins was set to all detected in the proteomics analysis. The list of enriched pathways and genes contained in each pathway were exported to Cytoscape (Paul Shannon et al., 2003). The EnrichmentMap app was used to generate a network of enriched pathways with the following parameters (pathway FDR p-value < 0.05; Jaccard index > 0.35) (Merico, Isserlin, Stueker, Emili, & Bader, 2010).

**Tissue sample preparation and Western Blotting Analysis**
Zebrafish muscle tissue (10-15mg) was placed in RIPA Lysis and Extraction Buffer (Thermo Fisher Scientific) with a cocktail of protease inhibitors and homogenized (2 X 15 seconds) using the Tissuemiser homogenizer (Thermofisher Scientific). Samples were separated on an SDS-PAGE and blotted onto polyvinylidene difluoride (PVDF) membranes. The membranes were blocked using 5% non-fat milk powder in 1X Tris Buffered Saline (Boston Bioproducts, MA) and 0.1% TWEEN® 20 (Sigma Aldrich, cat. no. P9416) (TBST) for 1 hour at room temperature and incubated with primary antibodies overnight at 4°C. The membranes were subsequently washed and incubated with polyclonal anti-mouse-IgG antibody conjugated to horseradish peroxidase. The antibody pairings and associated dilutions are:
Anti-KBTBD5 for KLHL40, 1:250 dilution (SAB2101198-100UL, Sigma Aldrich, PA); anti-$\alpha$-Tubulin,1:500 (ab18251, Abcam); anti-Sar1,1:100 (ab125871, Abcam); anti-Sec24d, 1:100 (14687, Cell Signaling technology); anti-Golga2, 1:100 (ab30637, Abcam); anti-FLAGM2 1: 250 (F1804, Sigma-Aldrich); anti-V5, 1:500 (R960-25, Thermofisher Scientific). Secondary antibodies were anti-rabbit 1:1000 (170-6515, Biorad) and anti-mouse, 1:1000 (170-6516, Biorad). The quantification of protein bands was performed using Image J.

**Immunofluorescence**
Zebrafish or human frozen skeletal muscle tissue were cryosectioned (8$\mu$m) and used for immunofluorescence as previously described (13) Antibodies used for immunofluorescence were anti-Sar1,1:100 (ab125871, Abcam); anti-RYR1,1:250 (R129, Sigma-Aldrich); anti-procollagen, 1:100 (MAB1912, Millipore Sigma); Integrin, 1:25 (clone8c8, DSHB). Secondary antibodies were anti-rabbit,1:250 (A11008, Thermofisher Scientific); anti-mouse,1:250 (A11005, Thermofisher Scientific).

**Sar1A overexpression in zebrafish**
Human *SAR1A* cDNA was subcloned from pDEST40-SAR1-V5-His6 plasmid (a gift from Richard Kahn; Addgene plasmid # 67451 ; http://n2t.net/addgene:67451 ; RRID:Addgene_67451) into pCSDest vector. mRNA was synthesized *in vitro* using mMessage kits (Ambion, Austin, TX, USA). 50–100 pg of mRNA was injected into embryos at the 1 cell stage.

**C2C12 Cell Culture Studies**

Coimmunoprecipitation of KLHL40 and Sar1A (from pDEST40-SAR1-V5-His6) was performed using the previously described method (12). To study the reciprocal interaction between KLHL40 and Sar1A, C2C12 cells were transfected with different amounts of KLHL40-pEZYFLAG and pDEST40-SAR1-V5-His6 plasmids. MG132 (10uM) was added at 40 hours post transfection, and cells were harvested at 48 hours post transfection. Cell lysates were prepared in RIPA buffer and proteins were analyzed by Western blot analysis.

**Electron microscopy**

Muscle tissue was dissected from juvenile and adult zebrafish, deskinned and fixed in formaldehyde–glutaraldehyde– picric acid in cacodylate buffer overnight at 4°C followed by osmication and uranyl acetate staining. Subsequently, tissue samples were dehydrated in a series of ethanol washes and finally embedded in Taab epon (Marivac Ltd., Nova Scotia, Canada). Ninety-five nanometer sections were cut with a Leica ultracut microtome, picked up on 100 m formvar-coated Cu grids and stained with 0.2% lead citrate. Sections were viewed and imaged by Joel 1200EX Transmission Electron Microscope (Electron Microscopy Core, Harvard Medical School).

**Expression and Purification of Sar1 and KLHL40 proteins**

Sar1A cDNA (addgene, #67451) or KLHL40 (WT or mutant cDNAs) were cloned into pDEST15 vector by gateway cloning. The Sar1-pDEST15 or KLHL40-pDEST15 vectors were transformed into BL21-Codon Plus (DE3) *E. coli* cells and cultured in LB media supplemented with 100 μg/mL ampicillin at a 1L scale. The cells were grown at 37℃ until $O.D_{600}$=0.4, and induced with 0.5 mM IPTG for 18h at 18℃. The harvested cells were resuspended in 25 mM HEPES (pH7.4), 130 mM NaCl, 20 mM $MgCl_2$, 1mM TCEP, 1mM PMSF, and 1 tablet of protease inhibitor cocktail (Pierce). Following lysis by French press, cell debris was pelleted by centrifuging at 15000 rpm for 40 min, and the soluble lysate was loaded on to glutathione-agarose resin (MCLAB) for affinity purification. The resin was washed with wash buffer containing 25 mM HEPES (pH7.4), 130 mM NaCl, 20 mM $MgCl_2$, 1mM TCEP, 0.1% Triton X-100, and then washed with additional wash buffer without Triton X-100. GST-tagged proteins were eluted with 50 mM reduced glutathione in 25 mM HEPES (pH7.4), 130 mM NaCl, 20 mM $MgCl_2$, 1mM TCEP, and dialyzed into 50 mM HEPES (pH7.4), 150 mM NaCl, 1 mM TCEP, 10% glycerol. The purified protein were concentrated to 5 mg/ml, flash-frozen, and stored at -80℃

***In vitro* ubiquitination assays**

The *in vitro* ubiquitination assays for Sar1A were conducted at 37°C in a total volume of 20 μL. The reaction mixture containing 5 mM ATP, 100 μM wild-type ubiquitin, 100 nM E1 protein, 2 μM E2 (UbcH5b), 0.38 μM CUL3-NEDD8-RBX2 (BostonBiochem, USA), 0.3 μM KLHL40 (wt or mutants) and 5 μM Sar1, with 40 mM Tris-HCl (pH 7.5), 50 mM NaCl, 0.5 mM TCEP and 5 mM $MgCl_2$ as the reaction buffer. Substrate Sar1 was preincubated with everything in the reaction mixture except E1 at 37°C for 20 min before E1 was added to the reaction system to initiate the reactions. Reactions were quenched at the indicated time points (0, 30 and 90 min) by adding SDS loading buffer containing reducing agent Dithiothreitol (DTT). The reaction samples were then resolved on SDS-PAGE gels and analyzed by either Colloidal Blue Staining kit (ThermoFisher Scientific, USA) or Western Blots. Assays were repeated on at least three independent occasions revealing results similar to the data presented in the figures.

**Western Blotting for *in vitro* ubiquitination assays**
After SDS-PAGE, the proteins were transferred to nitrocellulose membranes using an iBlot blotting system (ThermoFisher Scientific, USA). The membranes were then blocked with 5% BSA in PBST buffer for 1 h and then incubated with the anti-Sar1 antibody (1:500) at 4°C overnight. After this, the membranes were washed with PBST and probed with HRP-conjugated anti-Rabbit secondary antibody. The bands were detected by chemiluminescence using an Clarity Western ECL substrate (Bio-Rad, USA).

**Data analysis for *in vitro* ubiquitination assays**
To quantify the reaction rate of the Sar1 ubiquitination reactions, the mono-ubiquitinated Sar1 bands detected by Western Blot were quantified by densitometric analysis using Image J (version 1.53a). The relative ubiquitination rate of KLHL40 mutants group versus WT KLHL40 group were calculated from three biological repeats. The average values and standard deviations (presented as error bar) were calculated and shown in the figure. The statistical significance and $p$ values (or non-significant, n.s.) between groups were calculated using GraphPad Prism 9 using one-way ANOVA and reported in the figure.

**Table S10: Sequences of the sgRNA target sites and primers sequences (5'-3') to clone sgRNAs for creating *klhl40* zebrafish lines.**

| Gene | sgRNA Target site | Primers |
|---|---|---|
| *klhl40a-exon1* | GGACATCGAACCTGGCGTCA | F: TAGGACATCGAACCTGGCGTCA<br>R: AAACTGACGCCAGGTTCGATGT |
| *klhl40a-exon2* | GGCTGAGAACTCCATCTATG | F: TAGGCTGAGAACTCCATCTATG<br>R: AAACCATAGATGGAGTTCTCAG |
| *klhl40b-exon1* | GGACTGCATCAGGCTACG | F: TAGGACTGCATCAGGCTACGTC<br>R: AAACGACGTAGCCTGATGCAGT |
| *klhl40b-exon5* | GGACCGCAGTTCTCTCAGTC | F: TAGGACCGCAGTTCTCTCAGTC<br>R: AAACGACTGAGAGAACTGCGGT |
